# A uniquely leptin sensitive hypothalamic neuron population limits hyperphagia and weight gain in diet-induced obesity

**DOI:** 10.64898/2026.03.26.714161

**Authors:** Dylan M Belmont-Rausch, Benedicte Schultz Kapel, Abigail J. Tomlinson, Chao Ding, Allison M. Duensing, Bernd Coester, Cecilie Vad Mathiesen, Leonie Cabot, Jenny M. Brown, Charlotte Høy Kruse, Christoffer Clemmensen, Michael W. Schwartz, Henning Fenselau, Martin G. Myers, Tune H Pers

## Abstract

Despite widespread loss of leptin responsiveness in obesity, endogenous leptin continues to restrain feeding, yet the neural substrates that remain sensitive and mediate this effect are unknown. Combining spatial transcriptomics with single-nucleus RNA sequencing in mice with diet-induced obesity (DIO), we show that while most hypothalamic leptin receptor (*Lepr*) neurons minimally respond to elevated leptin, a single population defined by glucagon-like peptide-1 receptor (*Glp1r*) co-expression retains robust leptin sensitivity. These *Lepr/Glp1r* neurons project onto and restrain orexigenic *Agrp* neurons. *Lepr* deletion from *Lepr/Glp1r* neurons blocks the anorectic effect of exogenous leptin, reinstates hyperphagic responses normally suppressed in DIO, amplifies the obesogenic response to palatable diet, and unexpectedly attenuates hypothalamic microglial activation- a hallmark of DIO previously attributed to diet rather than leptin signaling. Hence, preserved leptin action through a single neuronal population governs downstream circuit activity to limit hyperphagia and weight gain during obesity.

## Introduction

Leptin is an adipocyte-derived hormone that signals through its receptor (LepR; encoded by *Lepr*) to communicate the repletion of body energy strores. Hypothalamic leptin action plays a central role to regulate energy balance by suppressing food intake and increasing energy expenditure ^1–4^. Obesity chronically elevates leptin and promotes a diminished anorectic response to exogenous leptin – a phenomenon broadly referred to as ‘leptin resistance’. This resistance is not complete, however, since interfering with leptin signaling exacerbates hyperphagia and weight gain in diet-induced obese (DIO) animals. Indeed, leptin deficiency causes obesity far beyond that observed in DIO ^5^. The specific neuronal populations that continue to mediate leptin action in DIO remain unclear, however, as do the roles for preserved leptin signaling in these neurons for the restraint of feeding.

The canonical targets of leptin action in the arcuate nucleus (ARC) – leptin-inhibited orexigenic agouti-related peptide (*Agrp*) neurons and leptin-activated anorexigenic pro-opiomelanocortin (*Pomc*) neurons ^6,7^ – both appear to become refractory to the direct effects of leptin in DIO ^8,9^. Previous work has shown that at least some *Lepr*-positive cell populations in the dorsomedial hypothalamus (DMH) continue to respond to leptin during obesity, however^10,11^. Moreover, even in the obese state, elevated leptin continues to restrain *Agrp* neuron activity through mechanisms that are not cell-intrinsic, suggesting that an upstream leptin-responsive population escapes the general loss of leptin sensitivity. Recent work identified a population of leptin-responsive DMH neurons that co-express glucagon-like peptide-1 receptor (*Glp1r*). While these GABAergic *Lepr/Glp1r* neurons are required for leptin’s normal restraint of body weight in chow-fed mice^12^ and project onto *Agrp* neurons^13^, their function in obesity is unknown. We hypothesize that these neurons maintain their leptin responsiveness to reduce hyperphagia in obesity by tonically inhibiting *Agrp* neurons.

Here, using spatial transcriptomics, single-nucleus RNA sequencing, and molecular genetic mouse models, we identify *Lepr/Glp1r* neurons as the principal neural population that exhibit increased leptin responses in DIO. We show that these neurons project onto and control gene expression in *Agrp* neurons during DIO. Removing leptin signaling from this single *Lepr/Glp1r* neuron population restores the sensitivity of *Agrp* neurons to orexigenic stimuli, amplifies the hyperphagic response to palatable diet, and attenuates hypothalamic microglial activation in DIO. These findings reframe our understanding of leptin action in obesity: rather than globally impairing leptin action, DIO promotes the progressive engagement of a specific neuronal population that limits hyperphagia and weight gain.

## Results

### A uniquely leptin-responsive hypothalamic neuronal population in obesity

To identify hypothalamic neurons that remain responsive to leptin in obesity, we combined phosphorylated STAT3 (pSTAT3) immunoreactivity (pSTAT3-IR; a readout of cellular LepR signaling), with spatial transcriptomics on the same tissue sections (**Fig. 1a**). Using a custom 300-gene panel, we profiled >312,000 cells from chow-fed lean and DIO mice (Supplementary Table 1, Extended Data Fig. 1a-c). We identified 17 distinct neuron populations **(Fig. 1b,c**; Extended Data Fig. 1d) that contained 7 previously-described *Lepr* populations, including ARC *Agrp*-, *Pomc*-, and *Tbx19*-expressing neurons, as well as a large GABAergic *Glp1r*-expressing population that spans the ventral DMH and caudal ARC ^14–16^. Label transfer confirmed correspondence of this latter population with previously described *Lepr/Glp1r* co-expressing mediobasal hypothalamic neurons ^14,17^ (Extended Data Fig. 1e).

**Figure 1.**
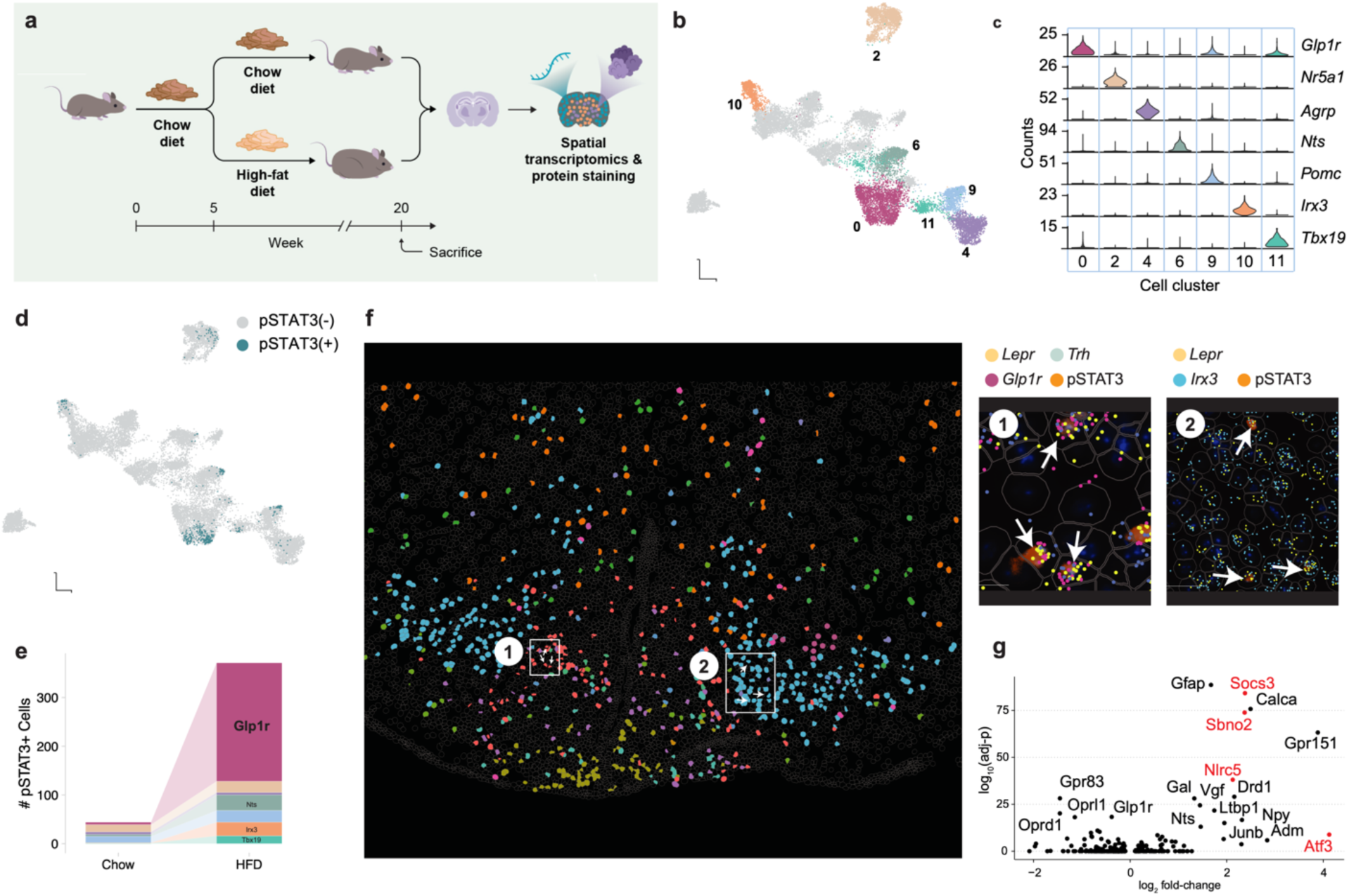
Elevated LepR signaling in hypothalamic *Lepr/Glp1r* neurons from DIO mice. **a,** Schematic of the combined Xenium spatial transcriptomics and pSTAT3 immunofluorescence workflow used to quantify leptin signaling across hypothalamic cell types in chow-fed and diet-induced obese (DIO) mice. **b**, UMAP embedding of neurons (17 populations; n=6 mice total) across all spatially profiled sections, colored by transcriptionally defined *Lepr* cell clusters. **c**, Marker gene expression across *Lepr* neuron subtypes, highlighting canonical cell type markers. **d**, UMAP embedding showing pSTAT3-positive *Lepr* neurons (green). **e,** Numbers of pSTAT3-positive cells across *Lepr* neuron subtypes in chow-fed and HFD-fed mice (n=3 mice per group; 2–3 sections per mouse). DIO increased pSTAT3 positivity nearly 10-fold overall (4.45±0.65% vs. 0.47±0.2%, *P*<0.001; generalized linear mixed-effects model with binomial distribution, animal as random effect). This was driven predominantly by *Lepr/Glp1r* neurons, which showed a >20-fold increase (21.1±1.1% vs. 0.33±0.2%, *P*<1×10⁻¹⁰) and accounted for >60% of all pSTAT3-positive *Lepr* neurons in DIO. *P*-values adjusted by Benjamini-Hochberg. **f**, Representative distribution of the each population of Lepr neurons across a single coronal section, color-coded as in b, c. Right panels show representative Xenium and immunofluorescence images showing spatial colocalization of *Lepr, Glp1r, Trh*, and pSTAT3 in the boxed areas of the larger panel. **g**, DIO-induced transcriptional changes in *Lepr/Glp1r* neurons, including increased expression of canonical leptin target genes (*Socs3, Nlrc5, Sbno2, Atf3;* red) and immediate-early genes (e.g., *Junb, Vgf*), indicating robust and sustained leptin signaling.

We detected pSTAT3-IR in the seven described *Lepr* neuron populations in the murine MBH (Methods, **Fig. 1d**). DIO increased the overall fraction of pSTAT3-IR *Lepr* neurons nearly 10-fold relative to chow controls (4.45±0.65% vs. 0.47±0.2%, P<0.001, n=3 mice/group; **Fig. 1e**; Extended Data Fig. 2a,b). This increase was not driven by pSTAT3-IR in canonical ARC leptin targets (i.e., *Agrp* or *Pomc* neurons) but was overwhelmingly concentrated in *Lepr/Glp1r* neurons, which accounted for more than 60% of all pSTAT3-positive neurons in DIO (>20-fold increase; **Fig. 1e**; Extended Data Fig. 2c,d).

DIO increased the expression of canonical LepR transcriptional targets (e.g., *Socs3, Nlrc5, Sbno2*) in *Lepr/Glp1r* neurons. Notably, the strong induction of *Socs3* – a well-characterized negative feedback regulator of LepR signaling –failed to terminate the transcriptional response of these cells to leptin, indicating that *Lepr/Glp1r* neurons remain leptin sensitive despite this feedback inhibition. Immediate early genes associated with neuronal activation (*Junb, Vgf*) were also increased by DIO in these neurons, suggesting increased neuronal activity in obesity (**Fig. 1g**). While a couple of other *Lepr* populations showed evidence of leptin sensing (increased pSTAT3) in DIO, the proportion of *Lepr/Glp1r* neurons that responded and magnitude of their response identified them as the principal responding population.

### *Lepr/Glp1r* neurons uniquely respond to a spectrum of nutritional states

The strong pSTAT3 response of *Lepr/Glp1r* neurons to DIO prompted us to ask whether these neurons also respond to leptin across a broader range of nutritional states known to differ in leptin concentrations. We therefore profiled more than 40,000 hypothalamic neuron transcriptomes across four conditions: fasting, refeeding, *ad libitum* chow-fed, and DIO (**Fig. 2a,b**; n=4-8 mice per group; Extended Data Fig. 3a–f). To identify which populations were most sensitive to these nutritional perturbations, we computed a transcriptional distance metric (*scDist*) that quantifies the shift in overall gene expression profile for each cell type relative to chow-fed baseline. Most populations responded robustly only to fasting, consistent with the established sensitivity of hypothalamic neurons to energy deficit. In contrast, *Lepr/Glp1r* neurons showed comparably strong responses to both fasting and DIO (**Fig. 2c**).

**Figure 2.**
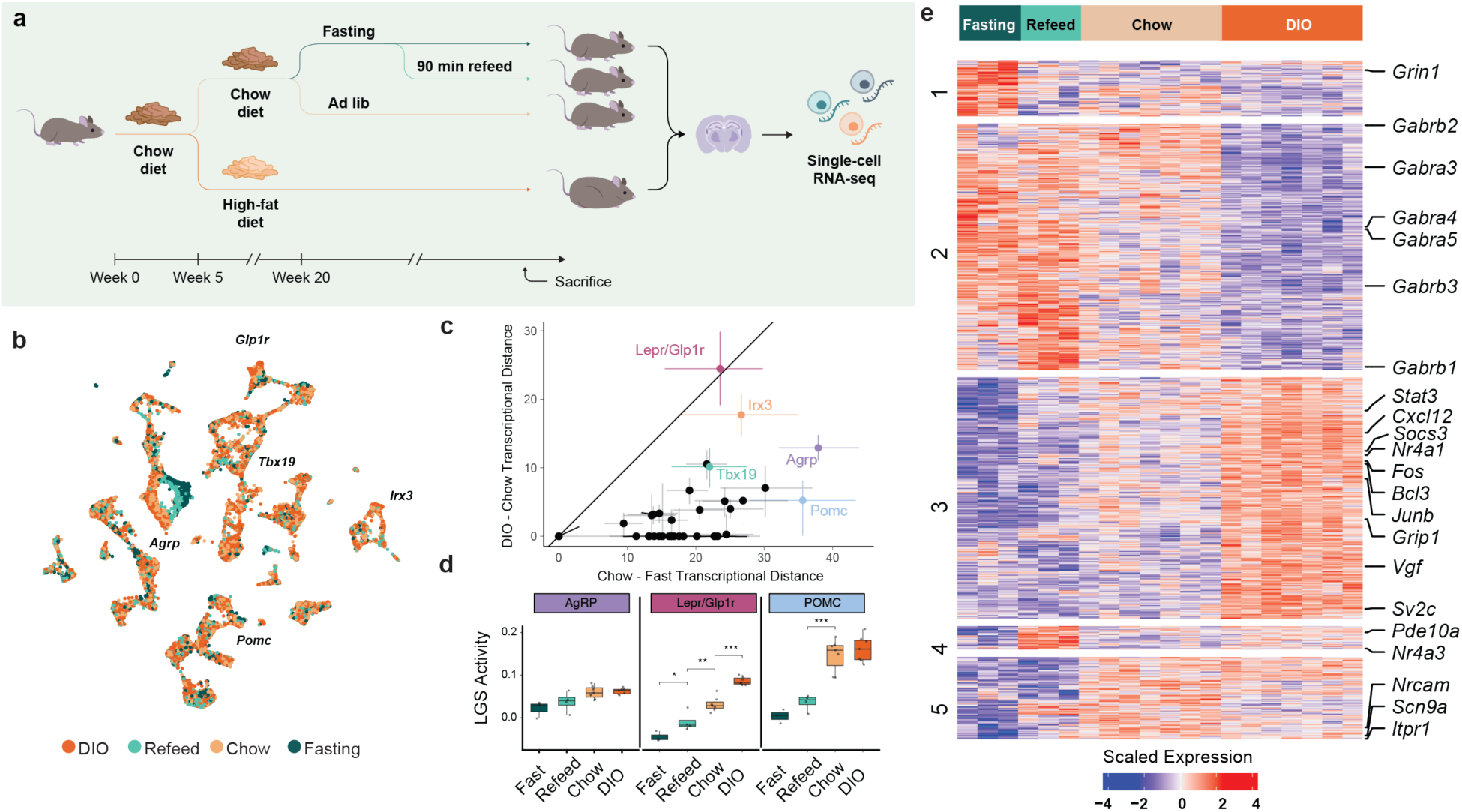
Uniquely elevated leptin signaling in *Lepr/Glp1r* neurons in DIO mice. **a**, Experimental design for snRNA-seq profiling of hypothalamic neurons across metabolic states: chow-fed (Chow; n=7), overnight-fasted (Fasting; n=3), fasted and refed (90 min) (Refeed; n=4), and DIO mice (HFD; n=8). **b**, UMAP embedding of MBH neurons (>40,000 nuclei; mean 12,401 transcripts per nucleus), with *Lepr*-expressing neurons labelled by subtype. **c**, Effects of nutritional manipulations on the total transcriptome of each neuronal population. Expression distance estimate in response to fasting (x-axis) and DIO (y-axis). Data for *Lepr* neurons is highlighted in cell-type specific colors. Solid diagonal line indicates matched effects between perturbations. **d**, Leptin gene signature (LGS) expression in for each nutritional manipulations across the indicated subsets of *Lepr* neurons; HFD (red), refeed (light green), chow (yellow), fasting (dark green)). **e**, Hierarchical clustering and grouping of differentially expressed genes identified in *Lepr/Glp1r* neurons. Neuronal activity genes and GABA receptor subunits highlighted.

Because these state-dependent responses could reflect multiple features of altered energy balance beyond leptin, we next asked which component was attributable to leptin signaling. To isolate the component of transcriptional change attributable to direct leptin signaling in each cell type, we derived a leptin gene signature (LGS), a composite score reflecting the coordinated expression of genes consistently regulated by leptin across three independent treatment datasets ^18–20^. This signature includes canonical LepR targets such as *Socs3, Etv6*, and *Atf3* (among others) (Extended Data Fig. 3g-i; Supplementary Table 2,3). Consistent with the pSTAT3 data (**Fig. 1**), HFD significantly increased LGS expression in *Lepr/Glp1r* (and *Irx3/5* neurons), but not other cell types. In contrast, fasting reduced LGS expression in *Lepr/Glp1r, Pomc*, and *Tbx19* neurons (**Fig. 2d**; Extended Data Fig. 3j). Thus, among the populations examined, *Lepr/Glp1r* neurons showed the clearest bidirectional leptin sensing response-strongly increasing in DIO and decreasing during fasting. Interestingly, *Agrp* neurons displayed a minimal LGS response across conditions, potentially consistent with a predominantly indirect effect of leptin on gene expression in these neurons.

This bidirectionality of the *Lepr/Glp1r* neuron transcriptional response extended beyond canonical leptin targets. In these cells, all genes that increased during fasting tended to decrease in DIO, and vice versa – a pattern of mirror-image regulation not seen in *Agrp* or *Pomc* neurons (Pearson’s r=−0.35, *P*=1.65×10⁻²⁸). This bidirectionality suggested that *Lepr/Glp1r* neurons actively track nutritional state across both energy deficit and surplus (Extended Data Fig. 3k).

Clustering of the 963 diet-regulated genes in *Lepr/Glp1r* neurons revealed a coherent transcriptional program (**Fig. 2e**). Genes associated with neuronal activation (*Fos, Vgf, Sv2c*) and cytokine signaling (*Stat3, Cxcl12*) increased progressively from fasting through chow to DIO. In contrast, GABA receptor subunits showed the opposite pattern, including downregulation of several GABAA subunits (*Gabra3, Gabra4, Gabra5*) and all three GABAB subunits (*Gabrb1–b3*) from DIO through chow to fasting (Supplementary Table 4-6). These data suggest that with increasing adiposity, *Lepr/Glp1r* neurons become more active and less sensitive to inhibitory input – a transcriptional profile consistent with their enhanced output onto downstream circuits.

To test whether leptin itself sufficed to induce this nutritional sensing program, we profiled single nuclei from lean mice treated with vehicle or leptin (3 mg/kg, IP) and harvested the MBH at 1, 3, 6, or 24 hours post-injection (n=4–6 per timepoint per group; >100,000 nuclei total; **Fig. 3a**; Extended Data Fig. 4a-d). *Lepr/Glp1r* neurons mounted the strongest response of any cell population (**Fig. 3c-d**; Extended Data Fig. 4e). Strikingly, leptin regulated 251 genes in *Lepr/Glp1r* neurons but only 28 in *Agrp* neurons (Supplementary Table 7; Extended Data Fig. 4f). Together with the minimal overlap of the leptin regulated genes from *Agrp* and *Lepr/Glp1r* neurons and the negligible LGS response in *Agrp* neurons, this near-complete absence of an overall transcriptional response to leptin in *Agrp* neurons (despite their expression of *Lepr*) is consistent with a predominantly indirect effect of leptin on *Agrp* neurons in vivo (*i.e.*, an effect mainly mediated through upstream populations such as *Lepr/Glp1r* neurons). Genes induced by exogenous leptin significantly overlapped with those upregulated in DIO in *Lepr/Glp1r* neurons (**Fig. 3e**), indicating that exogenous leptin administration can recapitulate a substantial component of the *Lepr/Glp1r* transcriptional program observed in DIO.

**Figure 3.**
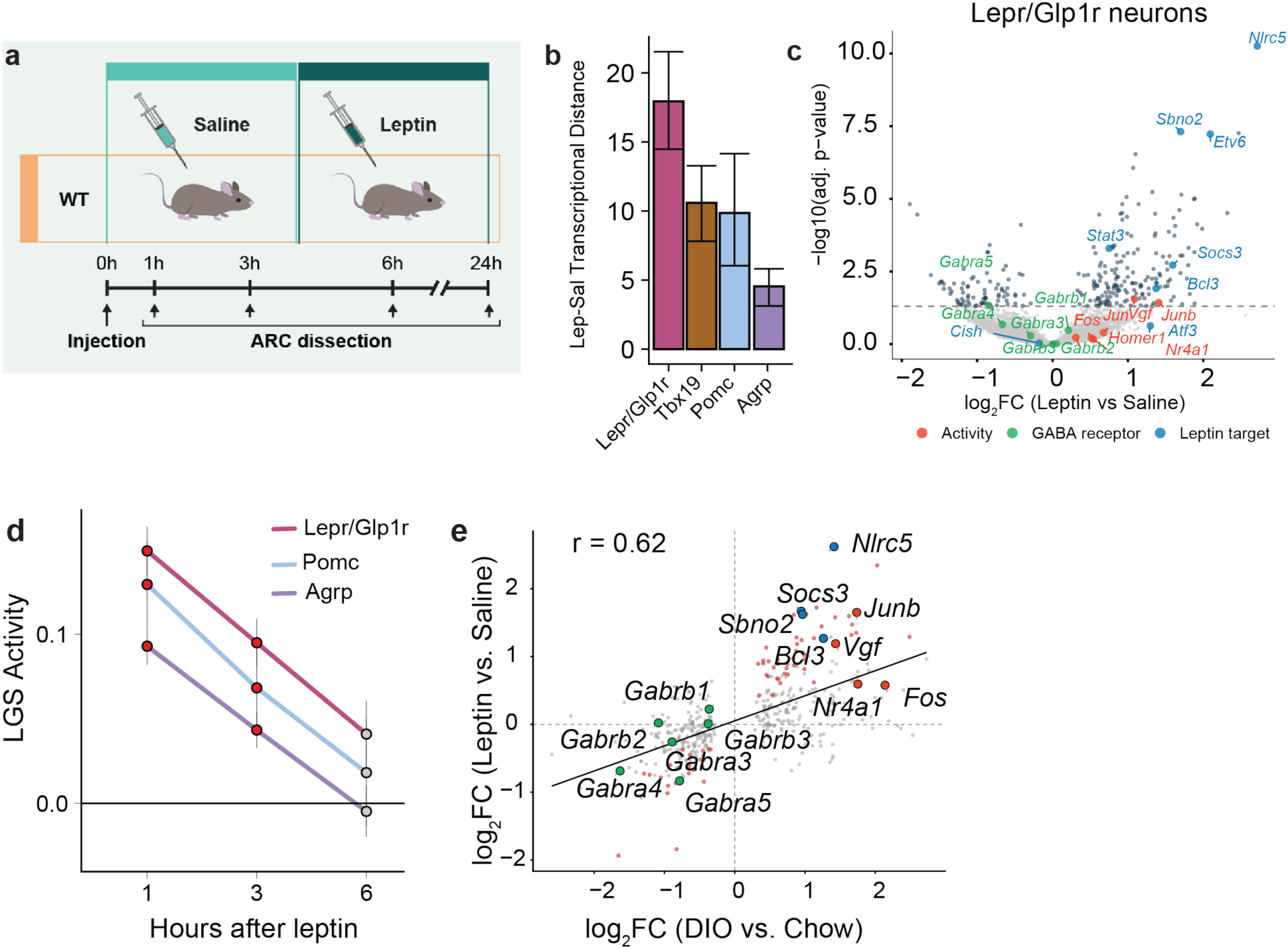
Acute leptin administration recapitulates the DIO transcriptional program in *Lepr/Glp1r* neurons. **a**, Experimental design of wild-type (WT) mice treated with 3 mg/kg of leptin of saline and sacrificed 1, 3, 6, or 24 hours after injection. MBH nuclei were collected for snRNA-seq. **b,** Transcriptional distance between cells from leptin treated and control mice. **c,** LGS changes across *Lepr/Glp1r*, *Pomc* and *Agrp* neurons**. d,** Overlap between genes induced by acute leptin treatment and those upregulated in DIO in *Lepr/Glp1r* neurons (93 shared genes; odds ratio=185.3, *P*=6.28×10⁻²⁴, Fisher’s exact test), indicating that direct leptin action recapitulates a substantial portion of the DIO transcriptional program.

### *Lepr/Glp1r* neurons in the ARC and DMH directly innervate *Agrp* neurons

Previous work established that *Lepr/Glp1r* neurons monosynaptically inhibit *Agrp* neurons^13^, but the relative contribution of this input among all LepR afferents has not been quantified.

Reanalysis of the published snRNA-seq data from rabies-traced *Agrp* neuron afferents revealed that *Lepr/Glp1r* neurons comprise approximately 80% of all *Lepr*-expressing GABAergic input to *Agrp* neurons (Extended Data Fig. 5a). These results identify *Lepr/Glp1r* neurons as the predominant inhibitory leptin relay onto *Agrp* neurons. While prior tracing only examined projections from ARC *Lepr/Glp1r* neurons^13,21^; our spatial data and previous findings^22^ show these cells also reside in the DMH. To determine whether *Lepr-* and *Glp1r-*expressing neurons in both locations project onto *Agrp* neurons, we performed two complementary monosynaptic rabies tracing experiments using *Npy^Flp^* on the *Lepr^Cre^*reporter background and *Agrp^Cre^* on the *Glp1r^Flp^*reporter background (**Fig. 4a,d**). In both strategies, starter cells were identified in the ARC (**Fig. 4b,e**) and rabies-labeled presynaptic neurons were traced to DMH *Lepr* and *Glp1r* neurons from ARC *Npy* and *Agrp* neurons, respectively (**Fig. 4c,f**). While the co-expression of *Lepr* and *Npy* in the ARC prevents the unequivocal demonstration that rabies-labelled ARC *Lepr* neurons lie upstream of *Npy^Flp^* neurons (as opposed to representing ARC *Lepr/Npy* neurons), the distribution of many traced *Lepr* cells in the dorsal lateral ARC (which contains few *Npy* neurons) suggests the presence of such afferents. *Agrp* neurons do not express *Glp1r*, however, and ARC *Glp1r* neurons were labelled with rabies retrogradely from *Agrp* neurons. Approximately 36% and 48% of rabies-traced presynaptic neurons in the ARC and DMH, respectively, were *Glp1r*-positive. Hence, *Lepr-* and *Glp1r*-expressing neurons in the DMH, as well as in the ARC, provide direct input onto *Agrp* neurons.

**Figure 4.**
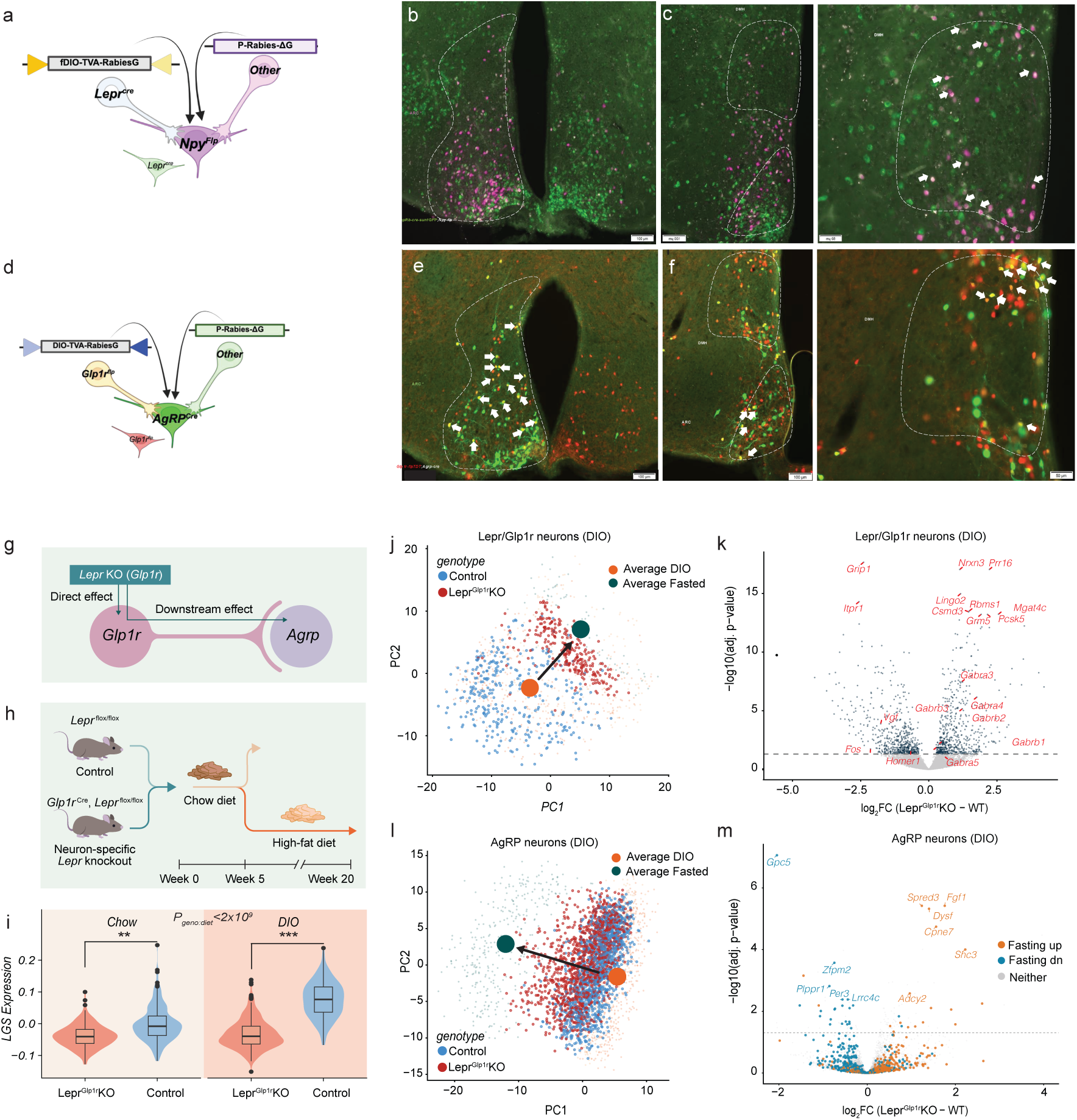
*Lepr/Glp1r* neurons project to *Agrp* neurons and leptin signaling via *Lepr/Glp1r* neurons controls *Agrp* neuron gene expression in obesity. **a**, Schematic of retrograde rabies tracing from ARC *Npy* neurons in *Lepr^Cre-sun1Gfp^;Npy^Flp^* mice. Starter cells (ARC) express TVA and rabies-G via *Npy^Flp^*; rabies-labeled cells are magenta, *Lepr*-expressing cells green, and co-labeled cells (*Lepr*-expressing afferents) show both signals. **b, c,** Representative images of tissue sections from experiments as in (a). *Indicates viral hit site. Afferent DMH *Lepr* cells indicated by white arrows. **d**, Schematic of retrograde rabies tracing from *Agrp* neurons in *Agrp^Cre;^Glp1r^Flp-TDT^* mice. Rabies-labeled cells are green, *Glp1r*-expressing cells red, and co-labeled cells (*Glp1r*-expressing afferents) show both signals. **e, f**, Representative images of tissue sections from experiments as in (d); *indicates viral hit site. Rabies-labeled *Glp1r* afferent cells indicated with white arrows. Scale bars: 200 µm (main images), 100 µm (insets). **g**, **h**, Experimental paradigm: *Lepr* was ablated in *Glp1r* expressing neurons and animals were fed chow diet (*Agrp*, n=7; *Glp1r*, n=6 *Lepr^KO^*and n=6 control) and either sacrificed at 4-5 weeks of age switched onto a HFD for 15 weeks (*Agrp* n=2; *Glp1r*, n=7 *Lepr^KO^* and n=6 control) until sacrifice. Mediobasal hypothalami were collected for snRNA-seq. **i,** Leptin gene signature (LGS) expression in *Lepr/Glp1r* neurons from for lean (Chow) or DIO Lepr^Glp1r^KO and control (WT) mice. LGS was significantly reduced in KO neurons (β=−0.087, *P*<1.0×10⁻⁹; linear mixed-effects model), with a significant genotype × diet interaction (*P*=2×10⁻⁹). **j,** PCA projection of *Lepr/Glp1r* neurons from DIO Lepr^Glp1r^KO (red) and Control (blue) mice onto the nutritional perturbation embedding. Centroids for DIO (orange) and fasted (teal) conditions shown as large circles. Lepr^Glp1r^KO neurons cluster with fasted wild-type neurons (PC1 permutation test, P=0.001), indicating LepR signaling is required to adopt the DIO transcriptional state. **k,** Volcano plot of differentially expressed genes in *Lepr/Glp1r* neurons (KO versus WT, DIO). Loss of *Lepr* abolished the DIO-associated induction of immediate early genes (*Fos*, log₂FC=−2.14, adj. *P*=0.018; *Vgf*, log₂FC=−1.76, adj. *P*=1.3×10⁻⁴; *Homer1*, log₂FC=−0.66, adj. *P*=0.027) and reversed the downregulation of GABA receptor subunits (*Gabra3, Gabra4, Gabra5, Gabrb1–b3*). Dashed line, adjusted *P*=0.05. See Supplementary Table 7 for full results. **l,** PCA projection of *Agrp* neurons from DIO Lepr^Glp1r^KO (red) and Control (blue) mice; centroids for DIO (orange) and fasted (teal) conditions shown as large circles, as in (**j**). *Agrp* neurons shift toward the fasted transcriptional state in Lepr^Glp1r^KO mice (PC1 permutation test, P=0.001), indicating propagation of the transcriptional effect from *Lepr/Glp1r* neurons. **m,** Volcano plot of differentially expressed genes in *Agrp* neurons (KO versus WT, DIO; 128 genes). Genes colored by their response to fasting: orange, fasting-upregulated; blue, fasting-downregulated; grey, neither. Fasting-upregulated genes were enriched among genes increased in KO (OR=8.56, *P*=1.05×10⁻⁸, Fisher’s exact test), and fasting-downregulated genes were enriched among decreased genes (OR=22.67, *P*=6.76×10⁻¹⁹), confirming a fasting-like transcriptional state despite obesity.

### Leptin action on *Lepr/Glp1r* neurons drives *Agrp* neuron transcriptional responses to obesity

Having established that *Lepr/Glp1r* neurons provide the predominant GABAergic LepR input to *Agrp* neurons, we asked whether cell-autonomous leptin signaling drives their adaptation to obesity and, consequently, mediates the response of *Agrp* neurons to DIO. We profiled neurons from mice lacking *Lepr* specifically in *Glp1r*-expressing neurons (Lepr^Glp1r^KO mice; **Fig. 4i**) on chow and after 15 weeks of HFD, when leptin is chronically elevated and the consequences of removing leptin signaling should be most apparent (35,538 cells; **Fig. 4j**; Extended Data Fig. 5b-d).

As expected, LGS expression was significantly reduced in *Lepr/Glp1r* neurons from Lepr^Glp1r^KO mice, with a significant genotype × diet interaction indicating that the effect was most pronounced under DIO conditions (**Fig. 4k**; Extended Data Fig. 5e). When projected onto the principal component embedding derived from the nutritional perturbation dataset (**Fig. 2b**), *Lepr/Glp1r* neurons from Lepr^Glp1r^KO mice clustered with those from fasted wild-type mice regardless of diet (**Fig. 4l**; PC1 shift permutation test, *P*=0.001), indicating that cell-autonomous LepR signaling is required for these neurons to adopt the DIO-associated transcriptional state. Loss of *Lepr* abolished the DIO-associated induction of immediate early genes (*Fos, Vgf, Homer1*) and reversed the downregulation of GABA receptor subunits in *Lepr/Glp1r* neurons (**Fig. 4m**; Supplementary Table 8-9).

The effects of loss of *Lepr* signaling in *Lepr/Glp1r* neurons propagated to *Agrp* neurons in obese animals. While *Agrp* neurons from chow-fed Lepr^Glp1r^KO mice were transcriptionally indistinguishable from controls, 128 genes were differentially expressed in *Agrp* neurons from DIO Lepr^Glp1r^KO mice (**Fig. 4n,o**; Supplementary Table 10-11). Notably, fasting-responsive genes were strongly enriched among the differentially expressed genes in *Agrp* neurons: those upregulated by fasting were overrepresented among genes increased in Lepr^Glp1r^KO mouse *Agrp* neurons, and those downregulated by fasting were overrepresented among genes with decreased expression in the DIO knockout animals (**Fig. 4o**). Projection of Lepr^Glp1r^KO Agrp neurons onto the diet PCA embedding confirmed a significant shift toward the fasting transcriptional state (PC1 shift permutation test, *P*=0.001; **Fig. 4n**). In other words, *Agrp* neurons in Lepr^Glp1r^KO mice maintain a fasting-like state despite obesity. Thus, direct leptin action on *Lepr/Glp1r* neurons is required to drive the response of *Agrp* neurons to DIO.

### Activation of *Lepr/Glp1r* neurons suppresses *Agrp* neuron-dependent feeding

If leptin acts through *Lepr/Glp1r* neurons to inhibit *Agrp* neurons, then direct activation of these neurons should suppress *Agrp*-driven feeding. To activate *Lepr/Glp1r* neurons selectively, we used an intersectional genetic approach requiring two recombinases (Cre and Dre) to restrict DREADD expression to neurons co-expressing *Glp1r* (via Cre) and *Trh* (via Dre)- markers whose combination identifies the *Lepr/Glp1r* population in the ARC and ventral DMH (Extended Data Fig. 6a,b). Using a within-subject crossover design, we found that chemogenetic activation of *Lepr/Glp1r* neurons suppressed dark-cycle food intake, reduced refeeding after an overnight fast, and blunted ghrelin-induced hyperphagia (**Fig. 5a-c**; Extended Data Fig. 6c). Thus, activation of *Lepr/Glp1r* neurons is sufficient to suppress *Agrp* neuron-dependent feeding processes.

**Figure 5.**
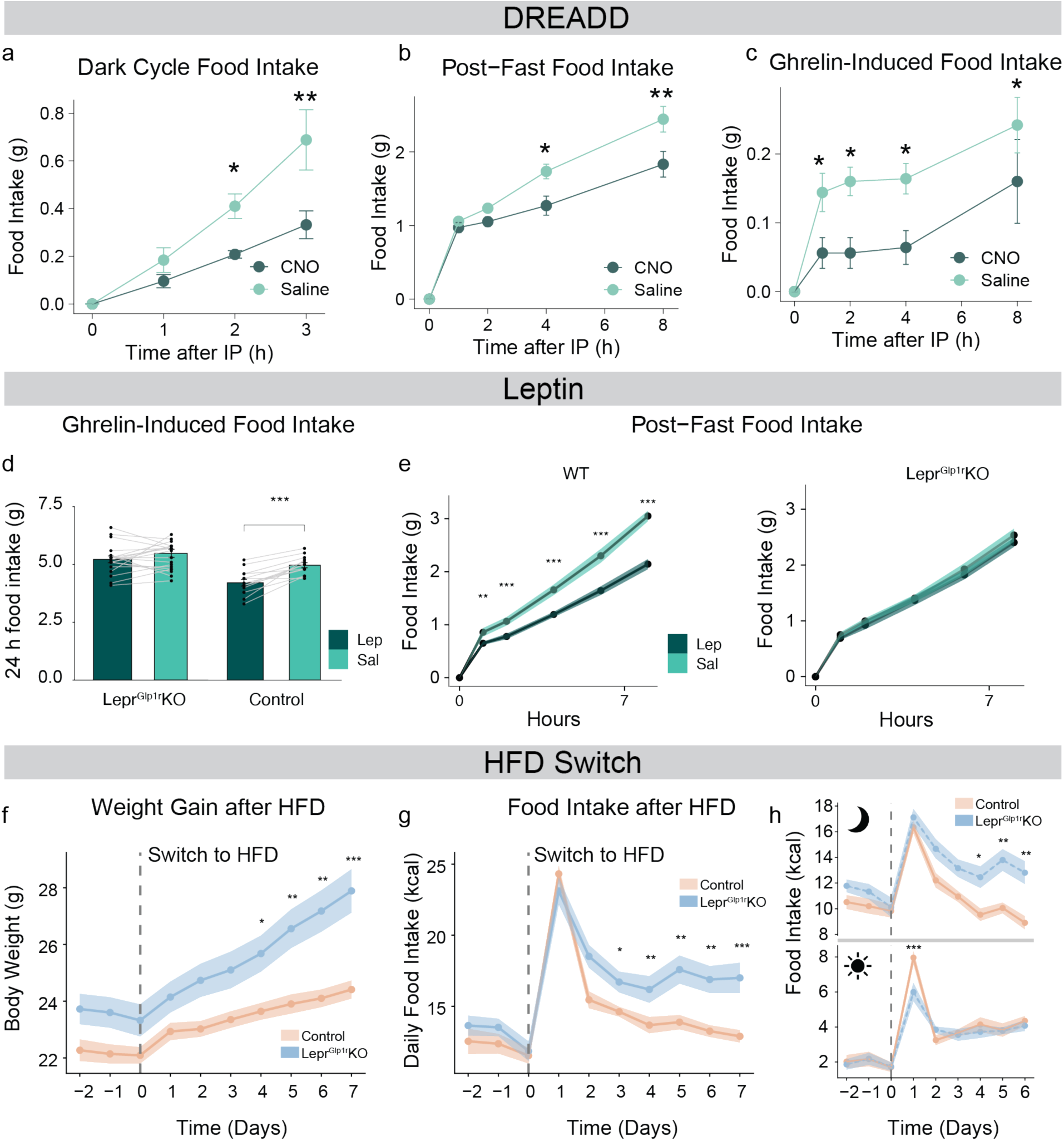
*Lepr/Glp1r* neurons restrain Agrp neuron-dependent and HFD-promoted hyperphagia. Mice containing activating DREADDs in Glp1r/Trh neurons were treated with saline or CNO at the onset of the dark cycle (a), during refeeding following an overnight fast (b), or prior to ghrelin treatment (c). **a**, Cumulative dark-cycle food intake at 0, 1, 2, and 3 hours following IP CNO or saline in a within-subject crossover design (n=5).. **b**, Cumulative post-fast food intake at 0, 1, 2, 4, 6, and 8 hours following IP CNO or saline (n=5). **c**, Cumulative food intake following IP ghrelin with CNO or saline pre-treatment at 0, 1, 2, 4, 6, and 8 hours (n=4). **d,** Effect of leptin (dark teal) versus saline (light teal) pre-treatment on ghrelin-induced 24-hour food intake in lean Lepr^Glp1r^KO (KO) and control (WT) mice. Lines connect within-subject measurements (crossover design). **e,** Cumulative post-fast food intake following leptin (dark teal) or saline (light teal) administration in WT (left) and KO (right) mice. **f, g**, Body weight (f) and daily food intake (g) before and 7 days after HFD exposure in Lepr^Glp1r^KO (n=9) and Control (n=10) mice. **h**, Food intake from (g) separated by dark (top) and light (bottom) cycle. The excess intake in Lepr^Glp1r^KO mice was concentrated in the dark cycle (genotype × time: χ²(9)=38.33, *P*=1.52×10⁻⁵). Dashed line indicates HFD switch. All panels: \**P*<0.05, **P<0.01, ***P<0.001. A-c, e-h: Plotted points represent mean values. Error bars (a-c) and shaded regions (e-h) denote SEM.

We next used Lepr^Glp1r^KO mice (Extended Data Fig. 6d) to ask whether *Lepr/Glp1r* neurons are required for exogenous leptin to suppress these behaviors. Exogenous leptin suppressed fasting-refeeding at every timepoint in wild-type mice but failed to reduce intake at any timepoint in Lepr^Glp1r^KO mice (**Fig. 5e**). Likewise, leptin failed to blunt ghrelin-induced feeding in Lepr^Glp1r^KO animals (**Fig. 5d**). Together, these findings identify *Lepr/Glp1r* neurons as the effector population through which exogenous leptin suppresses feeding responses known to be driven by *Agrp* neurons.

### Leptin signaling in *Lepr/Glp1r* neurons constrains hyperphagia during the onset of DIO

Having established that exogenous leptin requires *Lepr/Glp1r* neurons to suppress the hyperphagia that results from fasting or ghrelin treatment, we asked whether this mechanism operates during the physiological rise in leptin that accompanies weight gain. In wild-type mice a robust but transient hyperphagia accompanies the switch to HFD, typically peaking after 1–2 days and then normalizing over the following week as body weight is gained and leptin levels rise^23^. If leptin acts through *Lepr/Glp1r* neurons to restrain *Agrp* neuron driven feeding, then the progressive attenuation of this hyperphagia should require intact leptin signaling in these neurons.

Immediately following the switch to HFD, both Lepr^Glp1r^KO and control mice exhibited a comparable hyperphagia (**Fig. 5f**; Extended Data Fig. 6e). While food intake in control animals rapidly declined and returned to baseline by day 7, however, feeding remained elevated in Lepr^Glp1r^KO mice through the end of the study period (**Fig. 5g**). Consequently, Lepr^Glp1r^KO mice gained significantly more weight than controls (**Fig. 5f**). This prolonged excess food intake occurred predominantly during the dark (active) cycle, when *Agrp* neuron-driven feeding is highest (**Fig. 5h**), consistent with leptin acting via *Lepr/Glp1r* neurons to constrain *Agrp* neuron driven feeding.

### *Agrp* neuron desensitization in obesity requires leptin signaling in *Lepr/Glp1r* neurons

The reduced sensitivity of *Agrp* neurons to orexigenic stimuli, including fasting and ghrelin^24^, represents a hallmark of DIO. Because our data indicated that *Lepr/Glp1r* neurons tonically blunt the transcriptional response of *Agrp* neurons to DIO, we asked whether loss of leptin signaling in this population would restore *Agrp* neuron-driven responses in obese animals. We measured feeding responses to two canonical *Agrp* neuron-activating stimuli, fasting and ghrelin, in chow-fed and DIO Lepr^Glp1r^KO mice (Extended Data Fig. 7). Consistent with prior reports that obesity suppresses *Agrp* neuron responsiveness^24,25^, refeeding after an overnight fast was blunted in DIO wild-type animals, which consumed roughly half the calories of lean controls (**Fig. 6a**). In contrast, DIO Lepr^Glp1r^KO mice consumed similar amouns of food as lean controls, indicating that the DIO-associated suppression of post fast refeeding was largely abolished. Similarly, ghrelin failed to stimulate feeding in DIO wild-type animals but significantly increased intake in DIO Lepr^Glp1r^KO mice (**Fig. 6b**), consistent with restored *Agrp* neuron ghrelin sensitivity in these animals. Indeed, although baseline FOS-IR in the *Agrp* neuron-containing mediobasal ARC (mbARC) region did not differ between groups, ghrelin elicited increased mbARC FOS-IR in DIO Lepr^Glp1r^KO mice (**Fig. 6c**), consistent with restoration of *Agrp* neuron responsiveness. Together, these findings suggest that the normally reduced sensitivity of *Agrp* neurons in obesity reflects increased tonic inhibition from leptin-responsive *Lepr/Glp1r* neurons.

**Figure 6.**
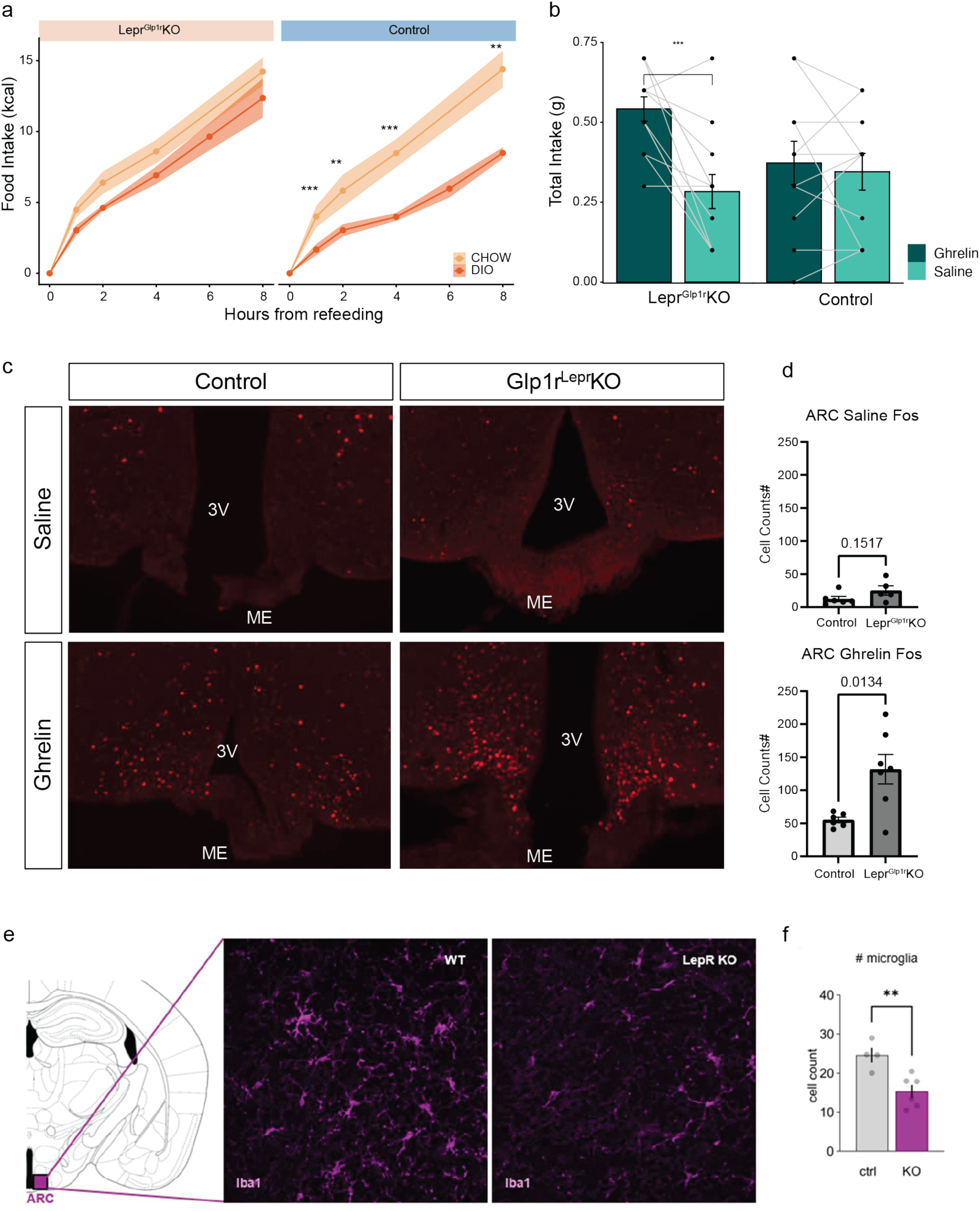
Leptin signaling in *Lepr/Glp1r* neurons mediates DIO-associated *Agrp* neuron suppression and hypothalamic microgliosis. **a**, Cumulative post-fast food intake (kcal) at 0, 1, 2, 4, and 8 hours in Control (WT, left) and Lepr^Glp1r^KO (right) mice on chow (light) or HFD (dark). \**P*<0.05, \*\*\**P*<0.001. Shaded regions denote SEM. **b**, Five-hour food intake following IP ghrelin (dark) or saline (light) in DIO WT (n=11) and Lepr^Glp1r^KO (n=12) mice. Lines connect within-subject measurements (crossover design). \*\**P*<0.01, error bars denote SEM. **c**, Representative FOS immunofluorescence in the mbARC of DIO Control and Lepr^Glp1r^KO mice following saline (top) or ghrelin (bottom) injection. 3V, third ventricle; ME, median eminence. **d**, Quantification of FOS-positive ARC neurons after saline (top; P=0.15) or ghrelin (bottom; *P*=0.013). Shown are mean-/+ SEM, along with individual data points. **e**, Left: schematic indicating the mbARC region sampled. Right: representative IBA1 immunofluorescence in DIO WT and Lepr^Glp1r^KO (KO) mice. **f**, Quantification of IBA1-positive microglia in the ARC of control (ctrl) and KO mice. Shown are mean-/+ SEM, along with individual data points.

### Loss of leptin signaling in *Lepr/Glp1r* neurons reduces hypothalamic neuroinflammation in obesity

Because Lepr^Glp1r^KO mice recapitulate several features of leptin deficiency, including preserved sensitivity to orexigenic stimuli despite obesity, we asked whether this phenotypic overlap might extend to hypothalamic neuroinflammation. Prior work has shown that, unlike DIO mice, obese leptin-deficient animals do not exhibit increased hypothalamic gliosis, suggesting that this hallmark of obesity depends in part on intact leptin signaling. Consistent with this idea, immunohistochemical staining for IBA1 in the ARC (**Fig. 6d**) revealed a significant reduction in microglial cell counts in Lepr^Glp1r^KO mice compared to wild-type controls (*P*<0.01), despite this greater food consumption and body weight. These findings suggest that obesity-associated MBH neuroinflammation is driven at least in part by leptin action on *Lepr/Glp1r* neurons.

Notably, each of the DIO-associated phenotypes altered by loss of leptin signaling in *Lepr/Glp1r* neurons – blunted *Agrp* neuron ghrelin sensitivity, attenuated fasting-refeeding responses, and hypothalamic microglial activation – is also absent in leptin-deficient *Lep^ob/ob^* mice despite comparable or greater obesity^24,26^. These parallels indicate that these hallmarks of DIO are not consequences of obesity *per se* but of sustained leptin action through *Lepr/Glp1r* neurons.

## Discussion

The concept of ‘leptin resistance’ rests on two observations: obesity associates with hyperleptinemia yet food intake remains normal or elevated, and exogenous leptin fails to reduce feeding and weight. Our findings reframe this phenomenon: rather than reflecting globally impaired leptin action, DIO represents a state in which rising leptin levels progressively engage a specific neuronal population – *Lepr/Glp1r* neurons – to limit hyperphagia and additional weight gain.

Our data also suggest that the pathogenesis of DIO involves multiple, distinct phases. Exposure to a palatable diet initially triggers a robust hyperphagia that appears largely independent of leptin signaling, consistent with the relatively low circulating leptin levels prior to the onset of substantial weight gain. This begins to change as body weight, fat mass, and leptin levels increase. While most *Lepr* neurons do not respond to these second phase changes in DIO, rising leptin levels continue to augment the activation of the uniquely leptin responsive *Lepr/Glp1r* neurons. This leads to the progressive inhibition of downstream *Agrp* neurons, which in turn likely contributes to the near normalization of food intake over the 7–10 days following the initiation of HFD feeding. In the final phase, sustained leptin action on *Lepr/Glp1r* neurons helps to establish a new, biologically defended body weight that, while higher than before, remains well below what would occur in the absence of leptin signaling.

The reduced overall responsiveness to exogenous leptin in DIO likely reflects two processes: the failure of hyperleptinemia to increase LepR signaling in most *Lepr* cell types plus the achievement of a ceiling effect for *Lepr/Glp1r* neurons. Not only might *Lepr/Glp1r* neurons reach maximal LepR signaling in DIO, but also *Agrp* neurons may be maximally inhibited (neuronal activity cannot go below zero). Notably, canonical feedback inhibitors do not explain why these neurons escape the general loss of sensitivity. SOCS3, a well-characterized terminator of LepR signaling^27^, is strongly induced in *Lepr/Glp1r* neurons during DIO yet fails to block their transcriptional response. Conversely, *Agrp* neurons show minimal *Socs3* induction yet respond poorly to elevated leptin. Nor do our transcriptomic data support previously proposed mechanisms such as inflammatory or ER-stress signaling in LepR neurons during DIO^28,29^. Instead, we propose that *Lepr/Glp1r* neurons operate across a wider dynamic range of leptin concentrations, enabling continued responsiveness as levels rise, while their sustained output onto *Agrp* neurons and other downstream targets accounts for the apparent unresponsiveness of those circuits to additional leptin.

Our data reinforce the emerging view that leptin’s effect on *Agrp* neurons is predominantly indirect, mediated through upstream GABAergic *Lepr/Glp1r* neurons^12,28^ that constitute the majority of inhibitory LepR input to this population ^13,30,31^. In DIO, this indirect pathway becomes the principal means by which leptin restrains *Agrp* neuron activity and limits feeding. Direct leptin action on *Agrp* neurons also contributes to the control of food intake and body weight, however, as mice with *Agrp* neuron *Lepr* deletion show modestly increased food intake and body weight on chow^32^. The more pronounced phenotype of Lepr^Glp1r^KO mice indicates that the indirect pathway plays the larger role. We note that both models involve germline *Lepr* deletion, howevr, and thus we cannot rule out developmental compensation.

Two additional consequences of sustained *Lepr/Glp1r* neuron activation merit comment. First, the well-documented insensitivity of *Agrp* neurons to ghrelin in DIO^24,25^ is reversed by removing leptin signaling from *Lepr/Glp1r* neurons, suggesting that ghrelin resistance in obesity is not cell-intrinsic to *Agrp* neurons but reflects extrinsic inhibition by this upstream population. Second, the attenuation of hypothalamic microgliosis in *Glp1r^Lepr-KO^* mice – mirroring the phenotype of leptin-deficient mice^26^ – indicates that obesity-associated neuroinflammation is driven by leptin signaling in *Lepr/Glp1r* neurons, rather than by adiposity *per se*.

In sum, selective preservation of leptin signaling in *Lepr/Glp1r* neurons links hyperleptinemia with the restraint of feeding in obesity, reframing what has been termed leptin resistance as a ceiling effect on a circuit that remains fully engaged. Several important questions follow from this model. Since leptin is unable to suppress intake of highly palatable food even in lean animals^33^, how do leptin-regulated circuits interact with neural pathways engaged by palatable foods? Our identification of persistently leptin-sensitive *Lepr/Glp1r* neurons as determinants of the response to this type of diet offers an avenue for investigating this question. Additional open questions include the molecular mechanism(s) underlying the failure of feedback inhibitors like SOCS3 to block LepR signaling during DIO links in *Lepr/Glp1r* neurons, the nature of the processes that restrain LepR signaling in other neurons, and the extent to which analogous circuit-specific leptin action persists in human obesity – where it might contribute to the strong association between obesity and hypertension through excessive leptin-mediated sympathetic outflow. Future work should also explore how leptin action on these neurons reshapes the response to other feeding-relevant signals, including cold exposure^34^, gut-derived satiety peptides^24^, and incretin agonists. Resolving these questions will be essential for determining whether the circuit-specific leptin action described here can be harnessed therapeutically.

## Acknowledgments

The Novo Nordisk Foundation Center for Basic Metabolic Research (CBMR) is an independent research center at the University of Copenhagen, partially funded by an unrestricted donation from the Novo Nordisk Foundation (Grant Agreement IDs: NNF23SA0084103, NNF18CC0034900). THP acknowledges funding from the Danish Council for Independent Research (Grant number 8045-00091B) and the National Institutes of Health (NIH) under grant R01 DK124238. DMR was fully supported by the Novo Nordisk Foundation Copenhagen Bioscience PhD Programme grant (NNF18CC0033668). MGM acknowledges support from NIH R01DK56731, the Michigan Diabetes Research Center (P30DK020572), and the Novo Nordisk Foundation/Center for Basic Metabolic Research. We would like to express our sincere gratitude to the Single-Cell Omics platform at the Novo Nordisk Foundation Center for Basic Metabolic Research for their contributions to this research project.

## Data and code availability

All data will, upon acceptance, be deposited in the Gene Expression Omnibus database and all code can be found at https://github.com/perslab/Rausch-2025.

## Author contributions

A.J.T, B.S.K, D.M.R, H.F., M.G.M., T.H.P designed the experiments, acquired and analyzed data. D.M.R performed all single-nucleus RNA sequencing data analysis. D.M.R wrote the first draft of the manuscript. A.J.T, B.S.K, D.M.R, M.G.M, M.W.S, and T.H.P wrote and edited the manuscript. All authors contributed to data acquisition, reviewed and edited the manuscript. M.G.M. and T.H.P. provided project oversight and acquired funding. T.H.P. is the guarantor of the manuscript.

## Declaration of generative AI and AI-assisted technologies in the writing process

During this work, the authors used ChatGPT and Claude to assist with R script preparation and to review sections of the manuscript for spelling and readability. After using these tools, the authors reviewed and edited the content as needed and take full responsibility for the content of the publication.

## Methods

### Mouse experiments

#### Animals and diets

C57BL/6N/J mice (Jackson Laboratory, #005304) were used as the wild-type group in the experiments. A *Glp1r*-specific knockout of *Lepr* (Lepr^Glp1r^KO mice) was produced by crossing the Lepr^flox^ mice with *Glp1r-ires-Cre* mice (Jackson Laboratory, #029283). For the DREADD experiments, *Glp1r-ires-Cre* (Williams et al., 2016); Jackson Laboratory, #029283) and *Trh-p2a-Dre* mice^35^ were utilized. *Npy^Flp^*, *Agrp^Cre^*, and the Flp- and Cre dependent reporter backgrounds (*Rosa26^FSF-tdTomato^* and *Rosa26^LSL-Sun1-sfGfp^* mice) were obtained from Jackson labs (Cat#: 030211, 012899, 032864, and 020139, respecitively) and bred in our colony. *Glp1r^Flp^* mice (a fusion of p2A-FlpO to the 3’ end of the endogenous *Glp1r* coding sequence) will be described elsewhere.

Throughout the studies, mice had *ad libitum* access to food (unless otherwise indicated) and water and were housed under specific pathogen-free conditions in temperature-controlled rooms with 75-80% humidity. Mice were maintained at a 12:12 h light cycle. All mice were fed the standard chow diets unless otherwise stated: the wild-type mice were fed Altromin 1310, Lepr^Glp1r^KO mice and their controls were fed Purina Diets 5LOD, and animals used for the DREADD experiment were fed ssniff®, V1154. The wild-type DIO mice were fed HFD with 58 kcal% fat and sucrose (Research Diets, D12331i), while DIO Lepr^Glp1r^KO mice and their controls were fed a 60 kcal% HFD (Research Diets, D12492). Only male mice were used for behavioral DREADD experiments. Experiments involving wild-type mice were conducted with approval from the Danish Animal Experiments Inspectorate under the permit numbers 2018-15-0201-01577 and 2020-15-0201-00767. Experiments with Lepr^Glp1r^KO mice and their controls were approved by the Institutional Animal Care and Use Committee at the University of Michigan (United States) under protocol number PRO00011066. The DREADD experiments were approved by local government authorities, Bezirksregierung Köln, Germany.

#### Xenium *in situ* gene expression analysis

Fresh frozen 10-12 μm tissue sections were placed on Xenium slides. The tissue was fixed and permeabilized according to the Xenium Fixation and Permeabilization Protocol (Demonstrated protocol CG000581). Probes were hybridized to the target RNA, ligated, and enzymatically amplified, generating multiple copies for each RNA target as described in Probe Hybridization, Ligation, and Amplification user guide (User guide CG000582). Xenium slides were then loaded for imaging and analysis on the Xenium Analyzer instrument, following the Decoding and Imaging user guide (User guide CG000584). Software analysis version 2.0.1.0 was used.

#### Immunostaining of brain tissue and image acquisition

Sections were brought up to room temperature (RT), air-dried, and slides were washed three times for 5 minutes each with phosphate-buffered saline (PBS) with Tween 20 (0.05%). Antigen retrieval was performed by incubating the slides in a solution of 0.3% NaOH and 1% H₂O₂ in PBS for 20 minutes. After two additional 5-minute washes with 0.02 M potassium phosphate buffered saline (KPBS), slides were incubated in 0.03% glycine in KPBS for 10 minutes, followed by two more 5-minute washes with KPBS. Blocking was carried out by incubating the tissues for 2 hours at RT in a blocking solution containing 5% normal goat serum (NGS) and 0.3% Triton X-100 in KPBS, adjusted to a total volume of 700 µL per slide. Phosphorylated STAT3 antibody staining was performed in KPBS with 1% NGS and 0.3% Triton X-100 with rabbit anti-pSTAT3 (Cell Signaling Technology, #9145; 1:200) overnight at 4°C. Slides were washed five times for 5 minutes each with 0.02 M KPBS. They were then incubated for 2 hours at RT with the secondary antibody, goat anti-rabbit AF594, diluted 1:200 in KPBS containing 0.4% Triton X-100. The samples were then counterstained with DAPI (0.5 mg/mL in KPBS) and then coverslipped using 80 µL of VECTASHIELD antifade mounting medium (Vector Laboratories, #H-1000) before being stored at 4°C. Images were acquired using a Axioscan Z1 scanning microscope with 20x magnification. Individual sections were segmented and analyzed in QuPath (version 0.5.0) ^36^. Nuclei were segmented on DAPI signal and classified as pSTAT3(+) after training an object classifier on individually identified positive cells.

#### Fasting/refeeding experiment

Wild-type mice were fed a chow diet until 5 weeks of age and subsequently divided into a HFD and chow fed group. At 20 weeks of age the chow fed group was split into a group being fasted overnight and a fed group. The fasting started at 6 p.m. The next morning the mice either stayed fasted or were refed chow diet between 6.00-8.00 a.m for 90 min before they were sacrificed together with the other study groups. In summary, this study consisted of the groups: a DIO (15 weeks HFD), a chow fed, a chow overnight fasted, and a chow refed group (90 min).

#### Leptin intraperitoneal injection experiments

For leptin intraperitoneal (IP) injections, 6 weeks old wild-type mice were acclimatized for 2 weeks before injection. Mouse recombinant leptin (R&D Systems, #498-OB) was reconstituted to 1 mg/ml in 20 mM Tris-HCl pH 8.0 (Invitrogen, #10434742). IP injections were conducted with 3 mg leptin/kg mouse or a corresponding volume of 20 mM Tris-HCl pH 8.0. All mice were group housed and leptin and vehicle injections were randomized across cages and litters. To allow dynamic, temporal assessment of the transcriptional leptin response in neurons, arcuate nuclei were collected from mouse brains 1, 3, 6 and 24 hours after leptin treatment. Mice were food deprived for 2 hours prior to injections in the 1 and 3 hour experiment.

#### Arcuate nucleus dissections

At sacrifice, brains were dissected and immediately placed in an ice-chilled coronal mouse brain slicer (Zivic Instruments, #BSMAS005-1, or ASI Instruments, RBM-2000C). A 1-1.5 mm thick coronal section was obtained between the rostral and caudal ends of Circle of Willis and placed in ice-cold PBS. Nuclei from the MBH of the left and right hemispheres were combined in a tube pre-cooled on dry ice and subsequently stored at −80 °C. All dissections were conducted between 8 and 12 a.m. to avoid transcriptional effects from circadian rhythm and feeding patterns.

#### DREADD stereotaxic surgical procedures

Mice were anesthetized with isoflurane and received an IP bolus of Buprenorphine (0.1 mg/kg bodyweight), and were put into a stereotaxic frame (David Kopf Instruments). A local anesthetic agent (Lidocaine) was applied to the skin, the skull surface was exposed through a skin incision, and a small drill hole was made. For virus injections, the AAV was unilaterally delivered through a pulled glass micropipette into the caudal ARC/ventral DMH (coordinates from bregma: AP: −1.5 mm, DV: −5.5 mm, ML: ± 0.4 mm; AP: −2.1 mm, DV: −5.5 mm, ML: ± 0.4 mm;). Before waking up, mice received analgesic treatment (subcutaneous injection of Meloxicam, 5 mg/kg) for pain relief and were carefully monitored to ensure regain of pre-surgery weight. All animals were allowed 4 weeks for virus expression before starting the experiment.

#### Food intake studies in *Glp1r^Cre^;Trh^Dre^* mice

AAV1/2-FLEX-FREX-hM3Dq (ETH Zurich, V892-1) was injected into the caudal ARC/ventral DMH of *Glp1r-ires-Cre;Trh-p2a-Dre* double transgenic mice at 8 weeks of age. Behavioral experiments were conducted four weeks post-injection. Food intake during the dark cycle was measured at 1, 2, and 3 hours after the onset of the dark cycle. Thirty minutes before the dark cycle, mice received an IP injection of either 3 mg/kg Clozapine N-oxide (CNO, Hello Bio, Cat# HB6149) or saline. For the fast-refeed experiment, mice were housed in fresh cages without food at the onset of the dark cycle and were subjected to a 16-hour fasting period. Food was reintroduced after 16 h of fasting and manually measured. CNO or saline was injected 30 minutes prior to food reintroduction. To investigate the orexigenic effects of ghrelin, CNO or saline was administered to fed mice at the onset of the light cycle. 30 minutes later, 1 mg/kg of ghrelin was injected IP immediately before food measurement.

#### Immunohistochemistry of *Glp1r^Cre^;Trh^Dre^* mice

For organ collection, mice were deeply anesthetized with ketamine and xylazine, and euthanized with transcardial perfusion of PBS, followed by 4% paraformaldehyde (PFA) in PBS (PFA-PBS). Brains were dissected, post-fixed at 4 °C in PFA-PBS for 16–22 h and then transferred to 20% sucrose in PBS. Brains were cut in 20 μm sections for immunostaining using a microtome. To cover the rostro-caudal axis of the brain region of interest, every fourth section was further processed (see below). The residual sections were collected in cryoprotectant and stored at −20 °C. Sections were blocked with 2% normal donkey serum in 0.4% Triton X-100 in PBS (NDS-PBST) for 1 h at RT and incubated with primary antibodies against HA (Rabbit HA Epitope Tag Antibody, ThermoFischer Scientific, 600-401-384) diluted 1:250 in NDS-PBST overnight at RT. Sections were washed with PBST and then incubated with secondary antibody anti-Rabbit IgG Alexa Fluor 555 (Life Technologies) diluted 1:1000 in PBS for 1 h at RT. After several washing steps with PBS, sections were mounted and counterstained with DAPI containing mounting medium (VECTASHIELD® Antifade Mounting Medium with DAPI, Cat# H-1200, Vector Laboratories). Brain sections were imaged by an olympus slidescanner (Olympus) with 40x magnification using the Cy3 filter.

#### Lepr^Glp1r^KO *in vivo* experiments

DIO male and female Lepr^Glp1r^KO (19 weeks on HFD) and age-matched chow-fed male and female Lepr^Glp1r^KO and appropriate littermate controls, were used to analyze the role of leptin-mediated control of ghrelin-induced feeding. Mice were allowed at least 3 days of recovery between experiments. To assess body composition, mice were placed in an EchoMRI (Echo Medical Systems). % mass was calculated in respect to individual body weight. Statistical analyses were performed using 2-Way ANOVA. Probability values <0.05 were considered statistically significant.

#### Lepr^Glp1r^KO ghrelin ± leptin feeding experiment

For the ghrelin ± leptin experiment, a cross-over study was performed in chow-fed Lepr^Glp1r^KO and control mice where order of exposure to leptin (Novo Nordisk, 5mg/kg) or an equivalent volume of 0.9% saline was randomized. 3 hours after the onset of the light cycle, food was removed, and mice were injected with either leptin or saline. 1 hour later, mice were injected with 1 mg/kg ghrelin IP (Hello Bio, HB2942) and food was returned. Food intake was measured at 24 hours post-refeed. Food intake was normalized to the average food intake of saline-treated *ad libitum* chow-fed control mice.

### Lepr^Glp1r^KO ghrelin chow to HFD feeding experiment

8 week old male and female Lepr^Glp1r^KO mice on a chow-fed diet were single housed, and twice daily food intake (following a 12:12h light cycle) and body weight were measured for 3 days to establish a metabolic baseline. Mice were then switched to HFD and caloric intake and body weight were followed for an additional 10 days. Body composition was measured prior to onset of the study, and 1 week and 2 weeks post-switch to HFD. Bodyweight was then followed weekly to determine onset of DIO prior to subsequent experiments.

#### Lepr^Glp1r^KO fast/refeed ± leptin experiment

A cross-over study was performed in chow-fed Lepr^Glp1r^KO and control mice where order of exposure to leptin (Novo Nordisk, 5mg/kg) or an equivalent volume of 0.9% saline was randomized. 3 days of baseline body weight and food intake were measured. On day −1, food was removed at the onset of the dark cycle and mice were injected with leptin or saline. The next morning, 3 hours after the onset of the light cycle, mice received a second injection. One hour later, food was returned (after a 16-hour fast), and food intake was measured at 24 hours post-refeed. Food intake was normalized to the average food intake of saline-treated *ad libitum* chow-fed control mice.

#### DIO Lepr^Glp1r^KO ghrelin feeding experiment

DIO male and female Lepr^Glp1r^KO and DIO control mice were injected with 0.9% saline (0.5 U/g) 4 hours after the onset of the light cycle. Food intake was measured after 5 hours. The following day, the animals were injected with 1 mg/kg ghrelin (IP) (Hello Bio, HB2942) and food intake was again measured after 5 hours. Food intake was normalized to the average food intake of saline-treated *ad libitum* HFD-fed control mice.

#### DIO Lepr^Glp1r^KO fast/refeed experiment

This study was performed in DIO male and female Lepr^Glp1r^KO or 8-week-old chow-fed male Lepr^Glp1r^KO mice, and appropriate diet-matched littermate controls. 3 days of baseline body weight and food intake were measured. On day −1, food was removed at the onset of the dark cycle. The next morning, food was returned after a 16-hour fast and food intake was measured at 0, 1, 2, 4, (6), 8, and 24 hours. Food intake was normalized to the average food intake of ad libitum HFD- or chow-fed control mice, respectively.

### Immunohistochemistry of Lepr^Glp1r^KO mice

For immunohistochemical analysis of neuronal activation after stimulation with ghrelin, *ad libitum* fed DIO male and female Lepr^Glp1r^KO and DIO control mice were injected i.p. with 1mg/kg ghrelin (Hello Bio, HB2942) or 0.9% saline (0.5 U/g). 90min post-injection, mice were anesthetized via isoflurane drop jar and euthanasia through transcardial perfusion of PBS followed by 10% formalin was performed. Brains were dissected and post-fixed in 10% formalin at RT for 4h prior and then transferred to 30% sucrose in PBS. Brains were mounted on a freezing microtome and sectioned at 35 μm. Hypothalamic sections were rinsed 3x in PBS prior to blocking in 3% normal donkey serum in 0.3% Triton-X 100 in PBS (PBST) for 1h at RT. Sections were then incubated overnight at RT in primary FOS (Cell Signaling Technology, 2250S 1:1000) or IBA1 () antibody. Sections were then washed 3x in PBS and incubated in secondary Donkey anti-Rabbit 647 (Invitrogen, A-31573 1:250) for 2h at RT. Following a final 3x PBS wash, sections were mounted and cover-slipped with Flouromount-G Mounting Medium (Southern Biotech, 0100-01). Imaging was performed on an Olympus BX53 microscope at 10x magnification using the cy5 filter.

### Single-cell sequencing

#### Nucleus isolation

Frozen arcuate nucleus samples were selected in pools of 4-12 randomized samples across developmental time and group. These samples were then recovered in an ice-cold Nuclei EZ lysis buffer (Sigma, #NUC101), lyzed and homogenized using a KIMBLE 2 ml tissue dounce with pastel B (Sigma, #D8938) 10 times. The homogenate was incubated for 5 min including pipette mixing 1-2 times. Cell debris and myelin were removed by filtering the homogenate through 40 µm mini cell strainers (Pluriselect, #43-10040) followed by centrifugation of the filtrate (1,000 rcf, 5 min, 4°C). The pellet was resuspended in a 1:1 solution of EZ lysis buffer and 50% OptiPrep™ iodixanol (Sigma, #D1556). Subsequently, the 25% iodixanol solution with nuclei was layered on a 29% OptiPrep™ iodixanol solution (Sigma, #D1556). Nuclei in the iodixanol gradient were centrifuged (14,000 rcf, 22 min, 4°C). The pellet was resuspended in nuclei buffer, PBS with 1% BSA, 2 mM Mg^2+^ and 40 U/µl Protector RNase inhibitor (Roche, #3335399001), and incubated for 15 min, before centrifugation (1,000 rcf, 10 min, 4°C). The final pellet was resuspended in 100 μl nuclei buffer for nuclei hashing.

#### Nuclei Hashing and sorting

To enable multiplexing of samples, a unique hashtag antibody was selected for each sample in a multiplexing pool. TotalSeq™ anti-Nuclear Pore Complex Proteins Hashtag antibody 0.5 µg/µl (Biologend, A0451-A0465, #AB_2861054 to #AB_2861068) was added to the 100 ul nuclei suspensions and incubated for 30 min. The nuclei were washed in nuclei buffer (1000 rcf, 10 min, 4°C) and resuspended in nuclei buffer with 0.5 µg/ml DAPI. Single DAPI^+^ nuclei were sorted on a SH800 cell sorter (Sony Biotechnology) using the 70 μm Sorting Chip (Sony Biotechnologies, #LE-C3207) directly into the Chromium Next GEM Single Cell 3’ v3 or v3.1 reverse transcriptase reagent (10x Genomics) and were used immediately for library preparation. We aimed to overload the Chromium chip with approximately 28,000 nuclei per reaction to optimize the capture rate of nuclei.

#### Library Preparation and RNA-Sequencing

Single nucleus RNA-sequencing libraries were generated using the Chromium Next GEM Single Cell 3’ kit, dual index, v3 or v3.1, with associated protocols (10x Genomics). The 10x Genomics protocols were followed up to cDNA amplification, where 1ul 0.2 uM hashtag oligo primers (see Supplementary Table 7) were added to the cDNA amplification mastermix. The cDNA amplification and the following SPRI bead clean-up was performed as described in the 10x protocol. However, the pellet and the supernatant were saved in separate 0.2 ml tubes. The 10x protocol was continued without changes for the pellet to generate libraries from endogenous RNA. To generate libraries of the hashtag oligos, 70 µl SPRI beads (Beckman Coulter, #B23317) were added to the supernatant. The solution was mixed 10 times and incubated at RT for 5 min. The tube was placed on a magnetic rack (10x Genomics) and the supernatant was discarded. The beads were washed twice on the magnet with 200 µl 80% ethanol, with 30 sec incubation per wash. After the second wash, the tube was briefly centrifuged and placed on the magnetic rack again. Any remaining ethanol was removed and the beads were air-dried for 2 min. The tube was removed from the magnet and 40 µl EB buffer (Qiagen, #1014609) was added immediately. The solution was pipette mixed thoroughly and incubated at RT for 2 min. The solution was placed on the magnetic rack again and 39 µl supernatant was transferred to a new tube and the DNA concentration was quantified using a Qubit Fluorometer (Invitrogen, #Q32851). To add sample indices to the hashtag oligo library, 50 µl Kapa HiFi HotStart ReadyMix 2X (Roche Diagnostics, #07958935001), 2.5 µl 10 µM SI PCR primer (see Supplementary Table 7) and 2.5 µl 10 µM Illumina TruSeq BioL_D70X_s (see Supplementary Table 7) were added to 20 ng hashtag oligo DNA. A library preparation PCR was performed (a. 2 min 98 °C, b. 20 sec 98 °C, c. 30 sec 64 °C, d. 20 sec 72 °C, repeated steps b.-d. 15 cycles in total, f. 5 min 72 °C, g. 4 °C hold). After the library preparation PCR, 120 ul SPRI was added to the sample, pipette mixed thoroughly and incubated for 5 min. The tube was placed on the magnetic rack and the supernatant was discarded. The beads were washed twice on the magnet with 200 µl 80% ethanol, with 30 sec incubation per wash. After the second wash, the tube was briefly centrifuged and placed on the magnetic rack again. Any remaining ethanol was removed and the beads. The tube was removed from the magnet and 40.5 µl EB buffer was added immediately, pipette mixed and incubated for 2 min. The tube was returned to the magnet, 39 µl supernatant was transferred to a new tube and the DNA concentration was estimated with the Tapestation High Sensitivity 1000 DNA kit (Agilent, #5067-5584) and Qubit Fluorometer (Invitrogen, #Q32851).

#### RNA sequencing

Single-cell libraries of endogenous RNA and hashtag oligos were sequenced on SP and S2 flow cells using the Illumina NovaSeq 6000 platform (Illumina) with Read1 length of 28 bp, Index1 length 8 bp and Read2 length of 94 bp, to a depth of approximately 40,000 reads per cell for the endogenous RNA library and 10,000 reads per cell for the hashtag oligo libraries.

### Data Processing & Analysis

#### Sequencing and Pre-Processing

BCL files were first demultiplexed into FASTQ files using bcl2fastq v.2.19.01. RNA reads were first aligned to an optimized mm10 transcriptomic reference gtf file using Cell Ranger version 7.0 including intronic sequences (10x Genomics). CellBender^37^ was used to mitigate the effects of contaminating ambient RNA on our analysis. Hash tag oligo (HTO) libraries were quantified with kb count, Kallisto Bustools version 0.24.4 ^38^. Quantified libraries were subsequently processed with Seurat ^39^. HTOs were normalized using the NormalizeData function with the normalization method set to CLR. A Gaussian mixture model with two components was fitted to each HTO distribution and a droplet was called to be positive for this HTO if it was predicted to belong to the cluster with the higher mean expression. Droplets positive for multiple HTO were classified as doublets. Inter HTO doublets were found using the recoverDoublets function from the scDblFinder package. Cells negative for HTO library as well as doublets were removed. Quality Control measures were calculated with AddPerCellQC(), Scuttle v1.4 (Bioconductor). Cells with too high or too low unique molecular identifiers (UMI) counts and intra- and intersample doublets were removed from the dataset. Genes expressed in less than 10 cells were removed.

#### Single-nucleus RNA-seq data analysis – initial processing

Downstream analysis was carried out in Seurat v4.3.0. For each 10X lane, cells with outlier UMI counts were identified and removed. Raw counts were normalized with the NormalizeData function, and principal component analysis (PCA) was performed with the RunPCA function using the top 3,000 variable genes as input. Subsequently, clustering was performed with the FindNeighbors and FindClusters functions using the top 30 PCs as input. Cells were split into neurons and non-neuronal cells based on the expression of known marker genes using cellAnnotator (Kharchenko Lab). For the neuronal and non-neuronal datasets from the diet perturbation, 10X lanes were integrated using the FindTransferAnchor and IntegrateData functions using 30 PCs. PCA and clustering were then run as described above for the neuronal and non-neuronal atlas separately using the top 40 PCs.

#### Label Transfer from Reference Single-Cell Datasets

To annotate cell types in our spatial and single-nucleus transcriptomics data, we performed label transfer from two independent reference datasets: a rabies-based hypothalamic cell atlas^17^ and a *Lepr*-expressing neuron dataset ^22^. Both reference datasets were preprocessed using standard Seurat workflows, including normalization, variable feature selection, scaling, and principal component analysis (PCA). Label transfer was performed using Seurat’s FindTransferAnchors and TransferData functions, utilizing the first 30 principal components. For each reference dataset, we computed prediction scores representing the confidence of cell type assignments. Cells were assigned a particular identity only if their prediction score exceeded 0.6; cells below this threshold were marked as unclassified. This conservative approach ensured high-confidence cell type annotations. We validated the consistency of transferred labels by examining their correspondence with our independently derived spatial transcriptomics clusters. The resulting annotations provided two complementary perspectives on cell type identity, leveraging both the comprehensive coverage of the AgRP neuron afferent rabies-based atlas and the specialized focus of the LepR neuron dataset.

#### UMAP Projection and Cell Type Label Transfer

To compare cell states across datasets while preserving the global structure of the reference data, we projected all datasets into the reference space defined by our nutritional perturbation atlas. We used the FindTransferAnchors() and MapQuery() functions to project query cells onto the reference UMAP embedding while simultaneously transferring cell type annotations.

#### Marker gene detection in Spatial Transcriptomics Data

To classify cells as positive or negative for specific marker genes in our spatial transcriptomics dataset we employed Gaussian mixture modeling. First, we performed centered log-ratio (CLR) normalization on the expression data to account for compositional bias. For each gene of interest, we analyzed the distribution of non-zero expression values using a two-component Gaussian mixture model implemented through the Mclust algorithm. This approach assumes that gene expression follows a bimodal distribution, with one component representing background/technical noise and the other representing genuine biological signal. Cells were classified as positive for a given gene if they belonged to the component with the higher mean expression level, as determined by the maximum likelihood estimates of the mixture model parameters. To validate the classification, we generated diagnostic plots showing the expression distribution overlaid with the fitted Gaussian components for each gene. This classification approach provided a systematic framework for identifying distinct cell populations in our spatial transcriptomics data while accounting for technical noise and probe diffusion.

#### Analysis of pSTAT3-Positive Cells

To assess differences in the proportion of pSTAT3-positive cells between dietary conditions across distinct spatial clusters, we employed a generalized linear mixed-effects model with a binomial distribution. The model included diet and cluster identity as fixed effects, with their interaction term, while accounting for animal-specific variation as a random effect. For each cluster, we computed total cell counts and the number of pSTAT3-positive cells. Estimated marginal means were calculated to test for diet-specific differences within each cluster, with p-values adjusted for multiple comparisons using the Benjamini-Hochberg method.

#### Spatial Differential Expression Analysis

To identify genes differentially expressed between dietary conditions, we performed statistical testing using the negative binomial test implemented in Seurat’s FindMarkers function. Raw counts were used for the analysis, with animal included as a latent variable to account for inter-individual variation. P-values were adjusted for multiple testing, and genes were considered differentially expressed at an adjusted p-value<0.05.

#### Identification of a Leptin Gene Signature Using Multi-Study Partial Least Squares Analysis

To identify genes regulated by leptin across multiple experimental contexts, we performed an integrative analysis of four independent transcriptomic datasets profiling leptin responses. These included TRAP-seq data from leptin-treated hypothalamic LepR neurons (GSE162603), RNA-seq from AgRP and POMC neurons after leptin treatment^40^, and RNA-seq data from nuclei of hypothalamic LepR neurons after leptin treatment (GSE112125).

For each RNA-seq dataset, we first performed quality control and normalization using DESeq2 v1.44.0. Raw count matrices were filtered to retain genes with at least 10 counts in a minimum of 2-3 samples (depending on the dataset). We then applied variance-stabilizing transformation (VST) to normalize the count data while accounting for the mean-variance relationship.

To integrate data across studies while accounting for batch effects and technical variation, we employed multi-study partial least squares discriminant analysis (MINT-sPLS-DA) using the mixOmics R package. Expression matrices from all four studies were merged based on common gene names. The model was constructed using two components and selected the top 100 most discriminative features (genes) per component. Data were scaled and near-zero variance features were removed prior to model fitting. Sample groups (leptin-treated vs. control) and study of origin were provided as categorical variables to guide the analysis.

#### LGS Scoring

LGS activity was computed using the AddModuleScore function in Seurat. For each leptin receptor-expressing neuronal population, we fit a linear mixed-effects models to test for differences in LGS activity across experimental conditions, while accounting for technical variation from cell hashing and sample preparation. The models included fixed effects for condition and random effects for sample ID and pool. Statistical comparisons between conditions were performed using estimated marginal means with Benjamini-Hochberg correction for multiple testing.

#### Quantification of perturbation effects

To quantify the effects of perturbations we used scDist (v1.1.0), a statistical framework designed to identify changes in gene expression while accounting for technical and biological covariates. Prior to analysis, gene expression data was processed using the SCTransform (Seurat). We used the scaled expression matrix consisting of the top 5,000 most variable genes as input and ran scDist with the following parameters: a minimum threshold of 5 counts per cell, treatment and hash-tag pool as fixed effects, and 10x lane identifier and sample as random effects. The analysis was conducted using 20 PCs.

### Pseudo Bulk Differential Expression Analysis

We performed pseudo bulk differential expression analysis using EdgeR (v4.0.16) and limma-voom (v3.58.1) to identify transcriptional changes between experimental conditions while accounting for technical and biological variation. Raw count data from single-cell RNA sequencing was aggregated into pseudobulk samples for each animal cluster combination using the Seurat2PB() function.

Samples with library sizes below 10,000 counts were excluded. We retained genes that met minimum expression thresholds of at least 1 count in a minimum number of samples and a total count of at least 5 across all samples. Library sizes were normalized using the trimmed mean of M-values (TMM) method.For differential expression testing, we used limma-voom. The design matrix incorporated fixed effects for experimental group and other technical batch effects (hash pool). Statistical significance was assessed using moderated t-statistics with Benjamini-Hochberg correction for multiple testing. Results were compiled across all analyzed cell types.

#### Principal Component Analysis Projection

To compare transcriptional states across datasets, we implemented a reference-based principal component analysis (PCA) projection approach. This method enabled direct comparison of cell states by projecting new data into a reference PCA space. For each cell type analyzed, we first identified shared genes between the reference and query datasets. The reference data was next log normalized and the top 5,000 most variable genes were identified for PCA.

The query dataset was independently normalized and scaled using the same set of variable features identified in the reference data. The scaled expression matrix from the query dataset was then projected into the PCA space defined by the reference data using matrix multiplication with the reference principal component loadings.

#### Diet Classification

To determine whether ablation of *Lepr* altered the transcriptional response to HFD, we developed an approach to classify cells based on their transcriptional state. We first projected neurons from genetic knockout models into the principal component space defined by control animals fed either chow or HFD diets. We next trained a binomial logistic regression model using the first five principal components from the reference dataset to predict diet condition (HFD vs chow). The model was then applied to the projected data from knockout animals to generate probabilistic predictions of diet state for each cell. These cell-level predictions were aggregated by animal to generate a mean prediction score for each biological replicate.

To assess the statistical significance of differences in prediction scores, we used a linear mixed-effects model. The model included fixed effects for dataset, genotype, and/or diet, and/or treatment. Post-hoc comparisons were performed using estimated marginal means with Tukey’s correction for multiple comparisons.

#### Statistical analysis

Statistical analyses were performed using R version 4.3.3 (R Core Team, 2018, https://www.rproject.org). Probability values<0.05 were considered statistically significant. Unless otherwise mentioned, the Benjamini-Hochberg procedure was used to adjust for multiple tests.

## Extended Data Figures

**Extended Data Figure 1.**
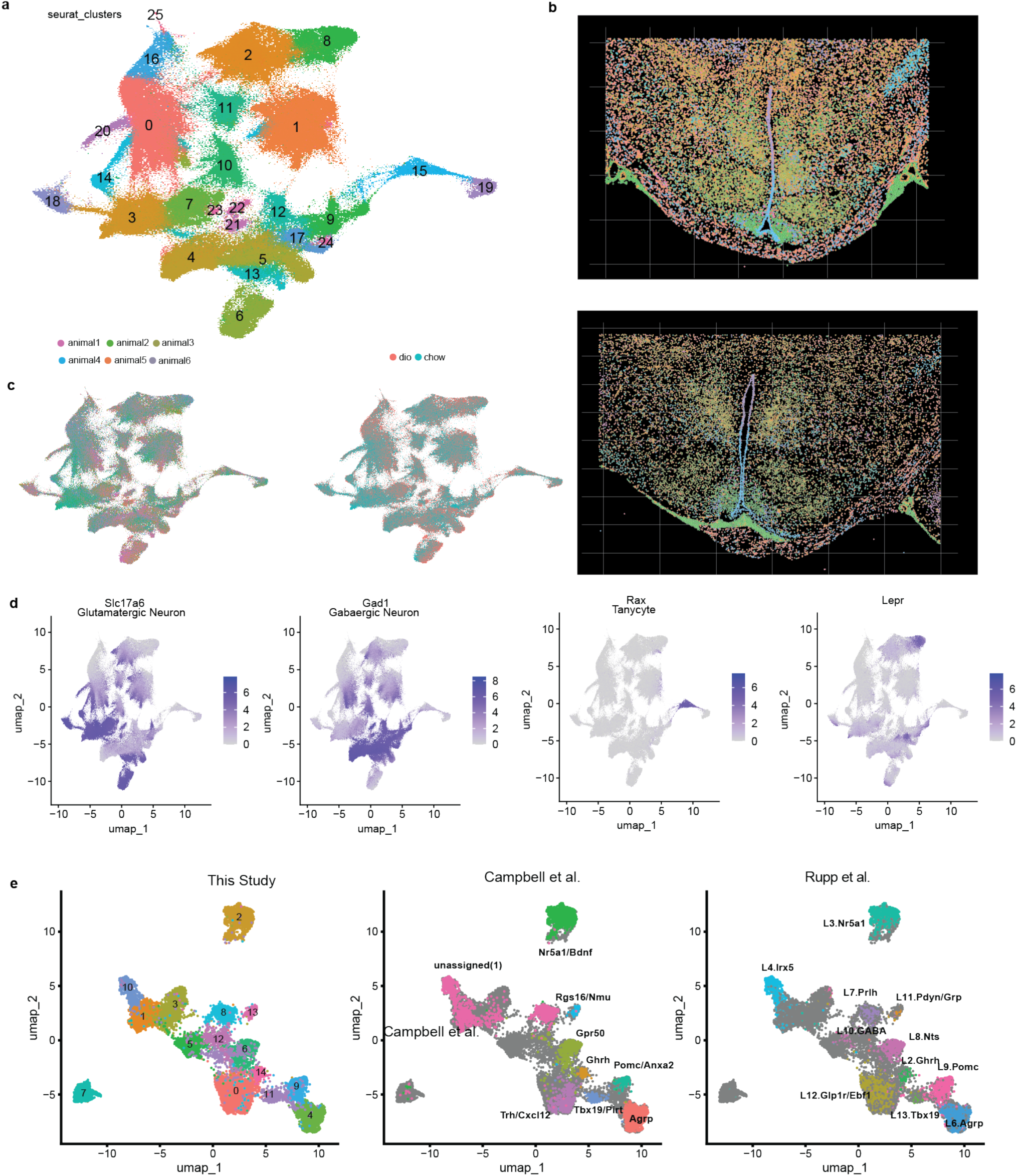
**a**, UMAP embedding of all cells from Xenium in situ sequencing annotated by seurat cluster numbers. Colors illustrate clusters. **b**, Representative images from Xenium in spatial context. Colors match clusters in panel a. Top image from Xenium slide 1 and bottom image from Xenium slide 2. **c**, UMAP embeddings of all cells from Xenium in situ sequencing annotated by animal (left panel) or by diet; 15 weeks HFD (DIO) and chow (right panel). **d**, UMAP embeddings of all cells from Xenium in situ sequencing colored by expression of *Slc17a6* (glutamatergic neurons), *Gad1* (GABAergic neurons), *Rax* (tanycytes) and *Lepr*. **e**, UMAP embedding of all neurons from Xenium in situ sequencing colored by seurat cluster (left panel), by Campbell et al. label transfer annotations (middle panel), and by Rupp et al. label transfer annotations (right panel).

**Extended Data Figure 2.**
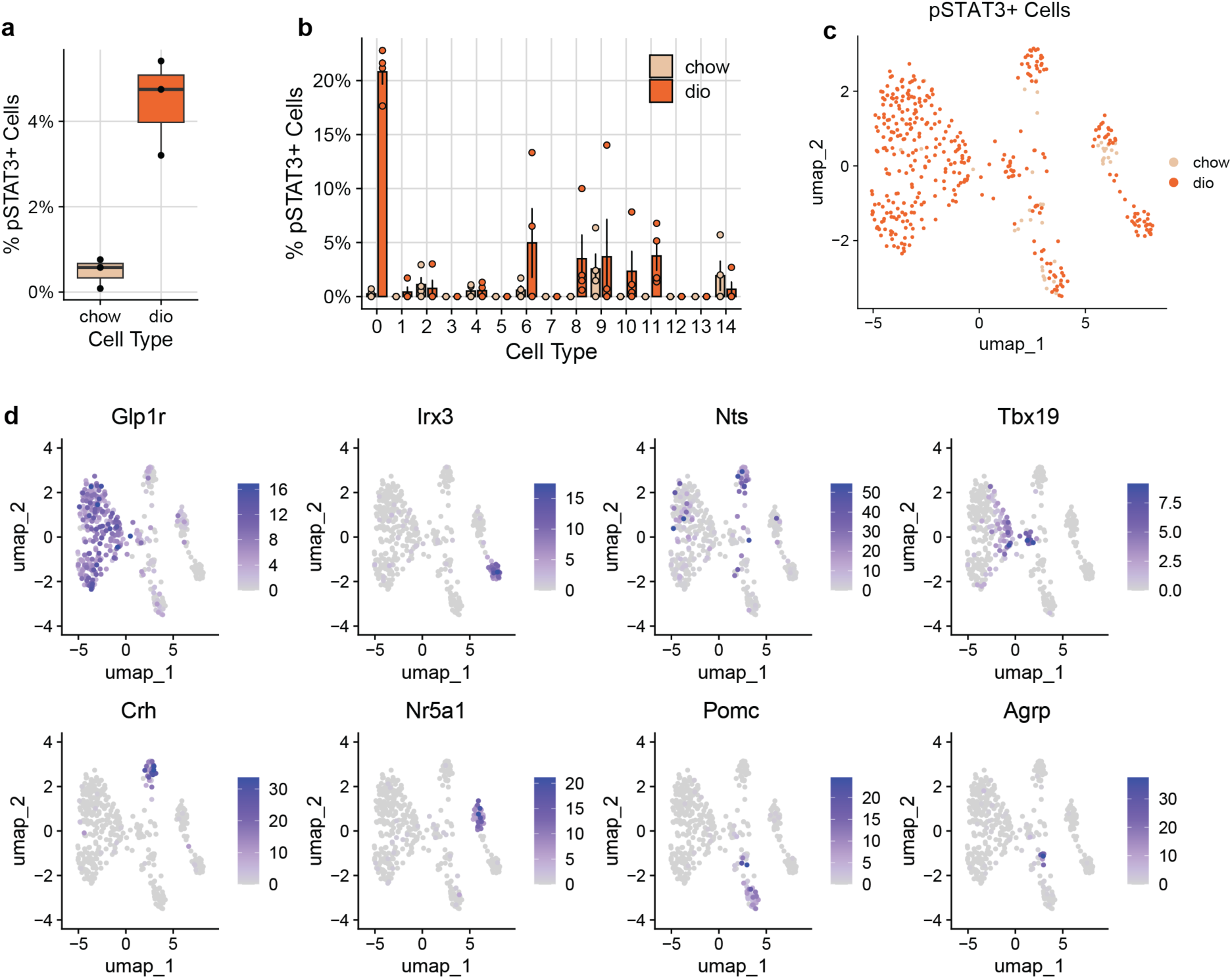
**a**, Quantification of pSTAT3-immunoreactive (pSTAT3-IR) cells in hypothalami of chow-fed and high-fat diet (HFD)-fed mice. **b**, Distribution of pSTAT3-IR cells across identified neuronal clusters in HFD-fed (red) and chow-fed (light orange) mice. **c**, UMAP visualization of Xenium in situ sequencing data from pSTAT3-IR cells, colored by diet condition (HFD: red; chow: light orange). **d**, UMAP visualization showing expression patterns of key neuronal marker genes (*Glp1r, Irx3, Nts, Tbx19, Crh, Nr5a1, Pomc,* and *Agrp*) in pSTAT3-IR cells.

**Extended Data Figure 3.**
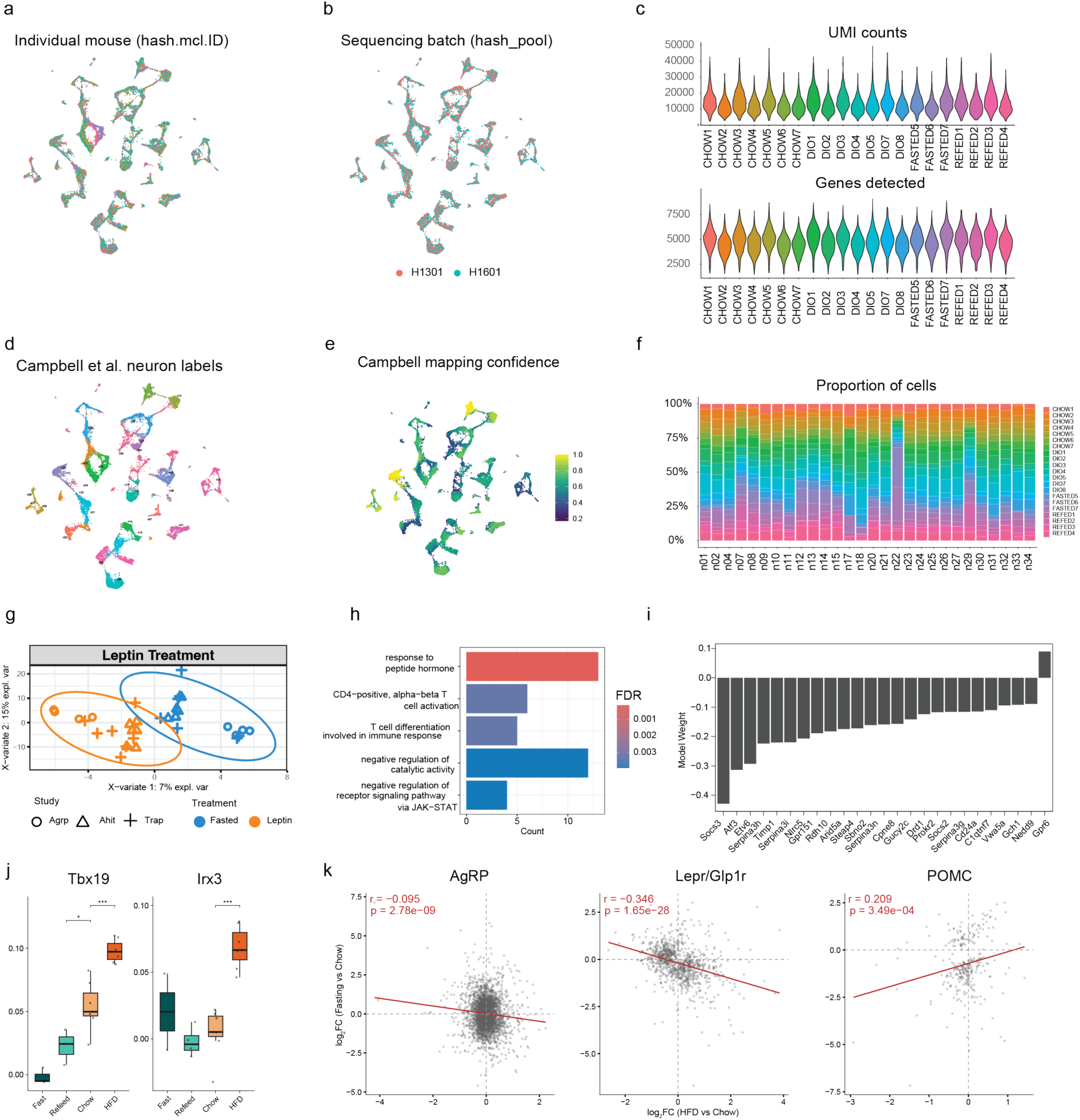
**a,b**, UMAP embedding of mediobasal hypothalamic neurons colored by individual mouse (a) and sequencing batch (b). **c**, Distribution of UMI counts (top) and genes detected (bottom) per cell across all samples (chow n=7, DIO n=8, fasted n=3, refed n=4). **d**, UMAP colored by predicted Campbell neuron subtype. **e**, Mapping confidence scores, with highest confidence in the Lepr/Glp1r population. **f**, Proportion of cells assigned to each neuronal cluster (n01-n34) across individual samples, showing consistent cluster composition across mice and conditions. **g**, MINT sPLS-DA projection of three independent leptin treatment transcriptomic datasets used to derive the leptin gene signature (LGS). Fasted/control (blue) and leptin-treated (orange) samples separate along the first two components. **h**, Gene ontology enrichment of LGS genes, highlighting response to peptide hormone and JAK-STAT signaling among the top terms. **i**, Model weights for individual LGS genes. **j**, LGS expression across nutritional states in Tbx19 and Irx3 neurons (***P<0.001). **k**, Correlation between fasting-induced (y-axis) and DIO-induced (x-axis) log₂ fold changes in AgRP, Lepr/Glp1r, and POMC neurons.

**Extended Data Figure 4.**
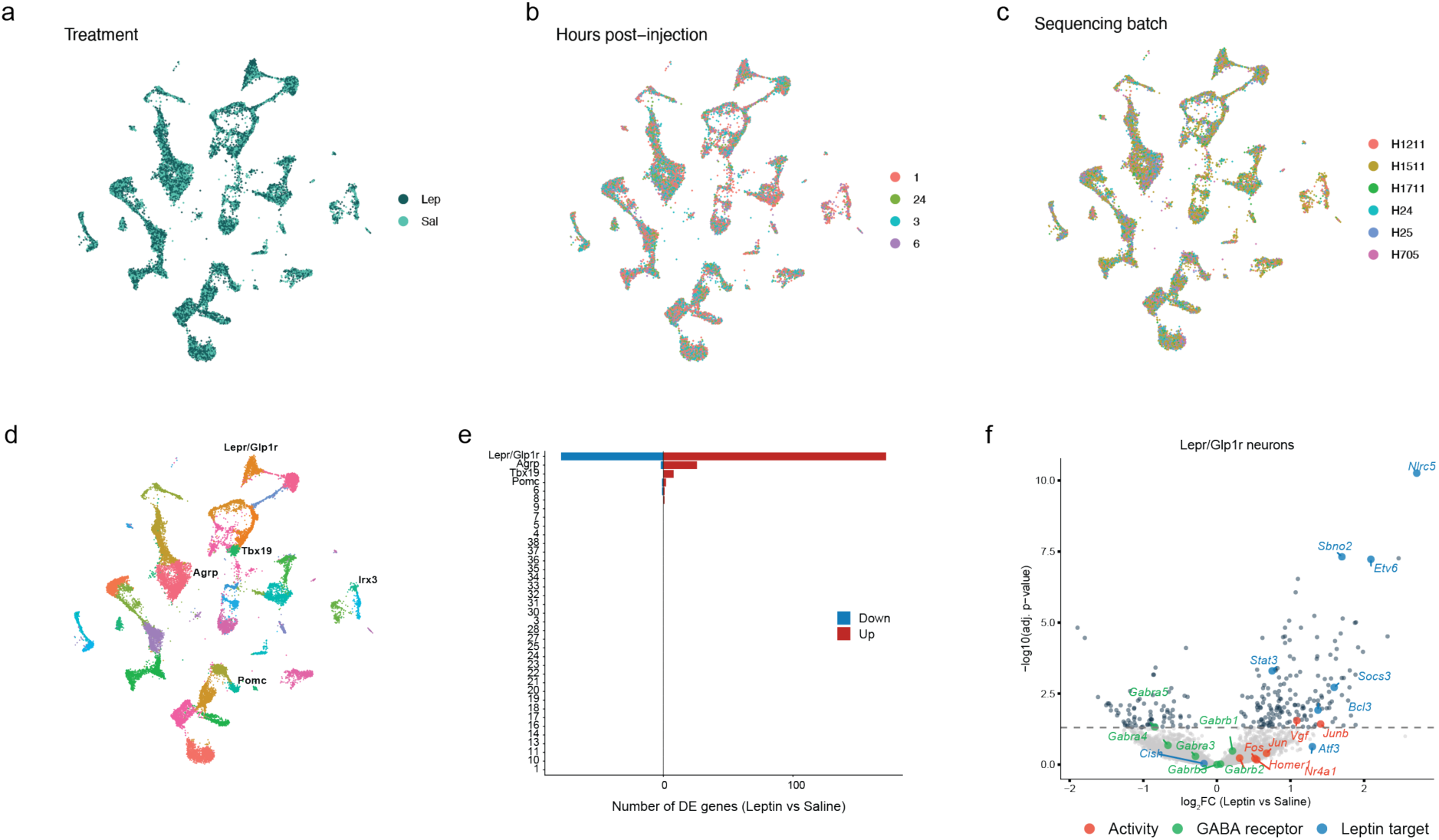
**a-c**, UMAP embedding of mediobasal hypothalamic neurons from lean mice treated with leptin (3 mg/kg) or saline and harvested at 1, 3, 6, or 24 hours post-injection (n=4-6 per timepoint per group), colored by treatment (a), hours post-injection (b), and sequencing batch (c). **d**, UMAP colored by neuronal cell type identity, with key Lepr-expressing populations labeled. **e**, Number of differentially expressed genes (leptin vs saline; adjusted P<0.05) per neuronal population. Red, upregulated; blue, downregulated. **f**, Volcano plot of leptin-induced differential expression in Lepr/Glp1r neurons. Highlighted genes include canonical leptin targets (blue), immediate early/neuronal activity genes (red), and GABA receptor subunits (green). Dashed line, adjusted *P*=0.05.

**Extended Data Figure 5.**
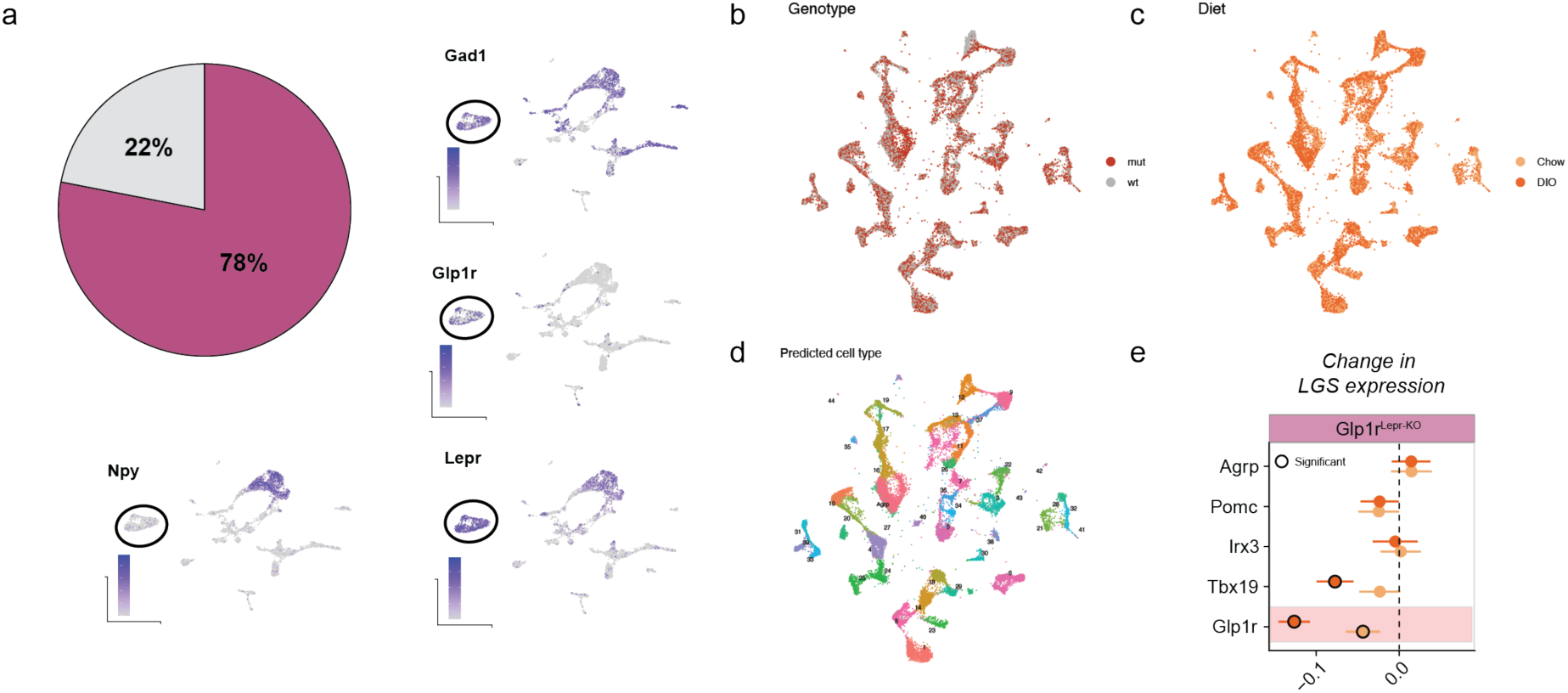
**a**, Lepr/Glp1r neurons constitute ∼78% of all Lepr-expressing GABAergic (Gad1+) input to AgRP (Npy+) neurons, based on reanalysis of published rabies-traced AgRP neuron afferents. Feature plots show expression of Gad1, Glp1r, Npy, and Lepr across traced populations; circled clusters indicate the Lepr/Glp1r population. **b,c**, UMAP embedding of mediobasal hypothalamic neurons from Glp1r^Lepr^KO and control mice (35,538 nuclei total), colored by genotype (b) and diet (c) (chow vs DIO; n=6-7 per group). **d**, UMAP colored by predicted cell type identity (41 clusters) based on label transfer from the nutritional perturbation reference atlas. **e**, Change in leptin gene signature (LGS) expression in Glp1r^Lepr^KO relative to control mice across *Lepr* neuron subtypes (open circles denote *P*<0.05; linear mixed-effects model).

**Extended Data Figure 6.**
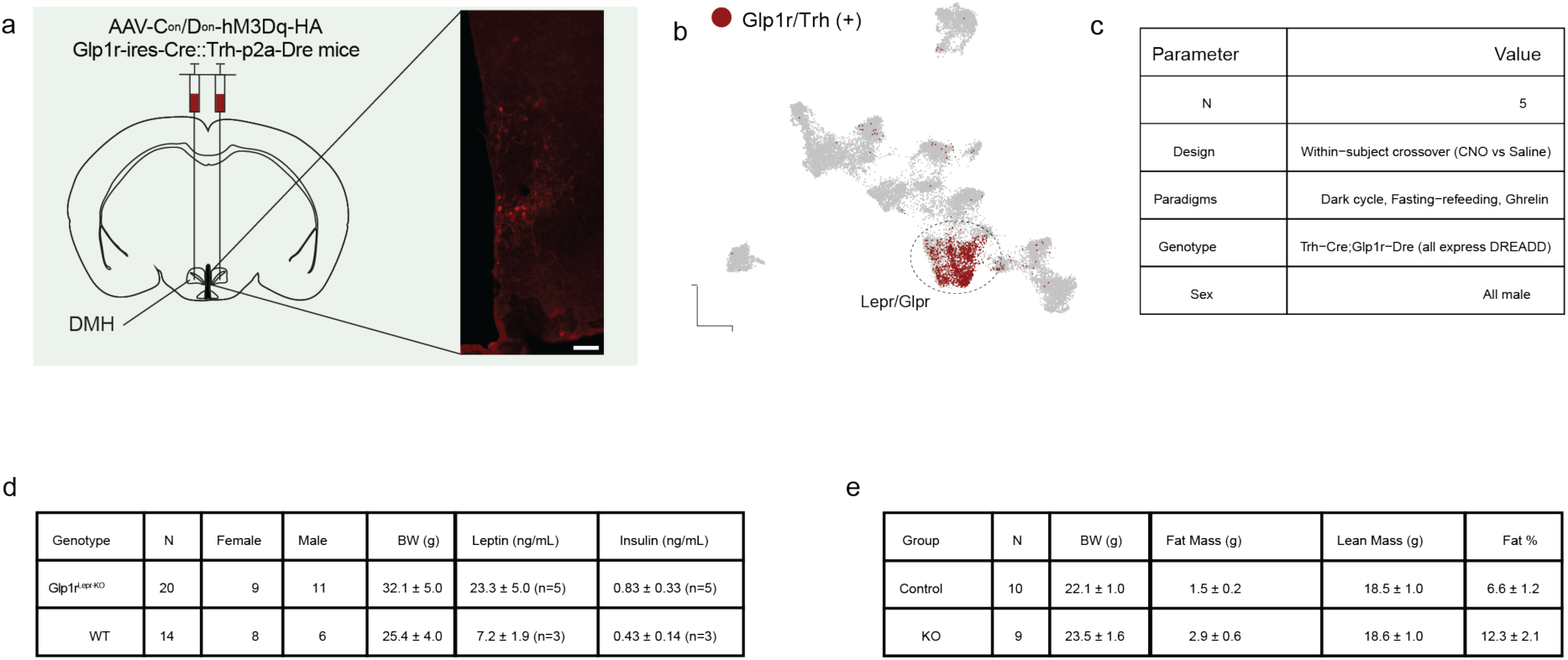
**a**, Schematic and representative histology of bilateral AAV-FLEX-FREX-hM3Dq injection into the caudal ARC/ventral DMH of Glp1r-ires-Cre;Trh-p2a-Dre mice. Scale bar: 200 µm. **b**, Spatial transcriptomics reference map showing the Lepr/Glp1r neuron population (red) targeted by the intersectional DREADD strategy. **c**, Summary of the DREADD cohort (main text Figure 4a-c). **d**, Baseline characteristics of chow-fed Glp1r^Lepr^KO (n=20) and control (n=14) mice used for leptin-dependent feeding suppression experiments (main text Figure 4d-e). **e**, Baseline chow-period body weight and DEXA body composition of Glp1r^Lepr^KO (n=9) and control (n=10) mice prior to HFD exposure (Figure 4f-h). Data are mean ± SD.

**Extended Data Figure 7.**
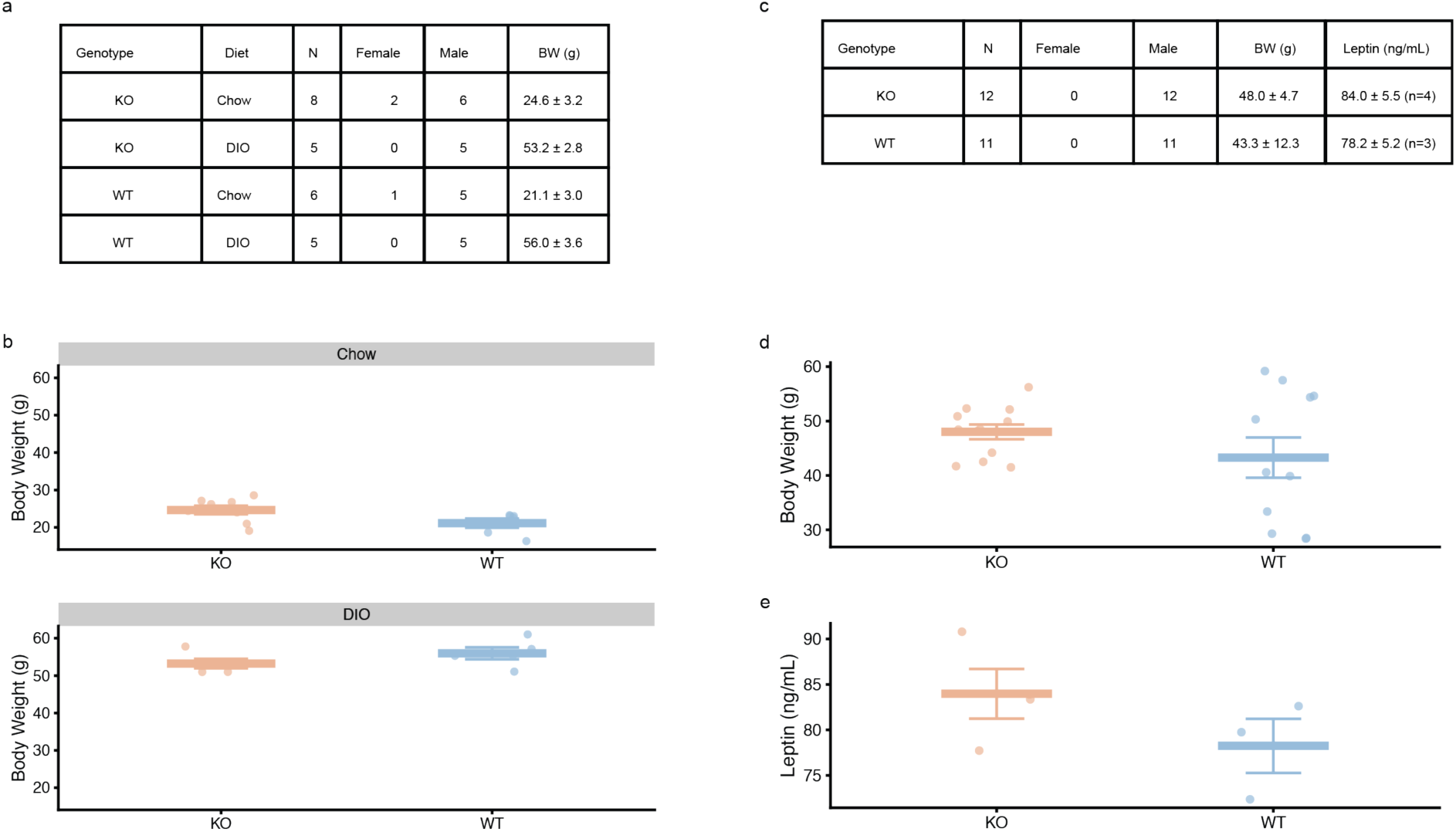
**a,b,** Baseline characteristics of Glp1r^Lepr^KO and control mice used in the chow vs DIO fasting-refeeding experiment (main text Figure 5a). Chow-fed and DIO cohorts are independent groups of animals. **c-e,** Baseline characteristics of DIO Glp1r^Lepr^KO (n=12) and control (n=11) mice used in the ghrelin feeding experiment. **c**, Summary table including plasma leptin for a subset of mice (KO n=4, WT n=3). **d**, Individual body weights and (**e**) plasma leptin by genotype, confirming hyperleptinemia in both genotypes. Data are mean ± SD (tables) or mean ± SEM (dot plots) with individual animals shown.

## Supplementary Table Legends

**Supplementary Table 1.** Custom 300-gene panel for Xenium spatial transcriptomics. Columns: gene, gene symbol; type, functional category (e.g., cell-type marker, leptin target, neuronal activation marker, body weight-associated receptor); ensembl_id, Ensembl gene identifier.

**Supplementary Table 2.** Leptin gene signature (LGS) genes and their weights. Genes were identified by multi-study partial least squares discriminant analysis (MINT-sPLS-DA) integrating three independent leptin treatment datasets. Columns: gene, gene symbol; value.var, loading weight reflecting each gene’s contribution to the discriminant component, with higher absolute values reflecting greater importance in distinguishing leptin-treated from control samples.

**Supplementary Table 3.** Gene ontology enrichment analysis of LGS genes. Columns: ONTOLOGY, ontology database (BP, biological process; MF, molecular function; CC, cellular component); ID, GO term identifier; Description, GO term name; GeneRatio, proportion of LGS genes annotated to the term; BgRatio, proportion of background genes annotated to the term; pvalue, unadjusted enrichment P-value; p.adjust, Benjamini-Hochberg adjusted P-value; qvalue, q-value for false discovery rate control.

**Supplementary Table 4.** Diet-regulated differentially expressed genes across all Lepr neuron populations. Differential expression was assessed using edgeR likelihood ratio tests (F-test across diet conditions: chow, HFD, fasting, refeeding) on pseudobulk profiles. Columns: ct, cell type; gene, gene symbol; logFC.treatmentHFD/Refeed/Fast, log₂ fold-change relative to chow for each condition; logCPM, average log₂ counts per million; F, F-statistic; PValue, unadjusted P-value; FDR, Benjamini-Hochberg false discovery rate.

**Supplementary Table 5.** Diet-regulated gene clusters in Lepr/Glp1r neurons. 963 differentially expressed genes (FDR < 0.05) were grouped into 5 clusters by hierarchical clustering of expression patterns across diet conditions. Columns: cluster, cluster assignment (1–5); gene, gene symbol.

**Supplementary Table 6.** Gene ontology enrichment analysis of diet-regulated gene clusters in Lepr/Glp1r neurons. Enrichment was performed separately for each gene cluster from Supplementary Table 5. Columns: Cluster, cluster number; name, cluster ID; ONTOLOGY, ontology database; ID, GO term identifier; Description, GO term name; GeneRatio, proportion of cluster genes annotated to the term; BgRatio, background ratio; pvalue, unadjusted enrichment P-value; p.adjust, Benjamini-Hochberg adjusted P-value; qvalue, q-value; geneID, gene symbols annotated to the term; Count, number of cluster genes in the term.

**Supplementary Table 7.** Differentially expressed genes between leptin-treated and vehicle-treated wild-type mice across Lepr neuron populations. Differential expression was performed using limma-voom on pseudobulk profiles for Lepr/Glp1r and Agrp neurons. Columns: gene, gene symbol; logFC, log₂ fold-change (leptin vs. vehicle); AveExpr, average log₂ expression across samples; t, moderated t-statistic; P.Value, unadjusted P-value; adj.P.Val, Benjamini-Hochberg adjusted P-value; B, log-odds of differential expression; CellType, neuron population (Lepr/Glp1r or Agrp).

**Supplementary Table 8.** Differentially expressed genes in Lepr/Glp1r neurons from chow-fed Lepr^Glp1r^KO vs. control mice. Differential expression was performed using limma-voom on pseudobulk profiles. Columns: gene, gene symbol; logFC, log₂ fold-change (KO vs. control); AveExpr, average log₂ expression; t, moderated t-statistic; P.Value, unadjusted P-value; adj.P.Val, Benjamini-Hochberg adjusted P-value; B, log-odds of differential expression; sig, significance flag (adj.P.Val < 0.05).

**Supplementary Table 9.** Differentially expressed genes in Lepr/Glp1r neurons from DIO Lepr^Glp1r^KO vs. control mice. Leptin reversed the diet-induced induction of immediate early genes and reversed the downregulation of GABA receptor subunits in Lepr/Glp1r neurons. Analysis as in Supplementary Table 8. Columns as in Supplementary Table 8.

**Supplementary Table 10.** Differentially expressed genes in Agrp neurons from chow-fed Lepr^Glp1r^KO vs. control mice. Agrp neurons from chow-fed Glp1r^LeprKO^ mice were largely transcriptionally indistinguishable from controls. Columns as in Supplementary Table 8.

**Supplementary Table 11.** Differentially expressed genes in Agrp neurons from DIO Lepr^Glp1r^KO vs. control mice. Loss of leptin signaling in Lepr/Glp1r neurons propagated to Agrp neurons in obese animals, with 128 differentially expressed genes. Fasting-responsive genes were strongly enriched among the differentially expressed genes, suggesting that Agrp neurons from DIO Lepr^Glp1r^KO mice adopt a fasting-like transcriptional state. Columns as in Supplementary Table 8.

